# A long-read human pangenome initiative for comprehensive interpretation of nuclear-embedded mitochondrial DNA

**DOI:** 10.64898/2026.02.26.708114

**Authors:** Lianting Fu, Jieyi Chen, Da Lian, Siyuan Du, Dongya Wu, Chentao Yang, Ziyi Wang, Hongyi Ma, Zhengtong Li, Nicole J. Lake, APG Consortium, Xiangyu Yang, Yongyong Shi, Guojie Zhang, Kaiyue Ma, Yafei Mao

## Abstract

Nuclear-embedded mitochondrial DNA segments (NUMT) preserve a record of ongoing mitochondrial-to-nuclear DNA transfer during evolution, with important implications for disease mechanisms and genome organization. Here, we developed a Pangenome Graph-based NUMT Detection (PG-NUMT) approach to improve NUMT detection sensitivity by 2.52-fold compared to short-read based approaches and comprehensively resolved seven concatenated mega-NUMTs with a maximum length of 127.7 kbp. We generated a high-resolution human NUMT map comprising 774 fixed, 280 polymorphic, and 123 pericentromeric NUMTs, alongside 74 superpopulation-stratified NUMTs. Both fixed and polymorphic NUMTs are hypermethylated and enriched within segmental duplications, whereas fixed NUMTs preferentially localize to intergenic regions and avoid transposable elements. Interestingly, NUMTs derived from the 3’-end of the mtDNA D-loop are less frequently fixed in human genomes and exhibit potential *cis*-regulatory activity in the nuclear genome. These observations suggest that selective pressures shape the genomic features of fixed NUMTs. Seven NUMTs associated with gene expression or alternative splicing were further identified, suggesting potential modulatory functions of common NUMTs. Using 20 complete non-human primate genomes, we identified variable NUMT insertion rates across primate lineages (1.5-18.7 insertions per million years), with particularly high rates in the *Pan* lineage. Notably, we uncovered two NUMT-derived variable number tandem repeats (VNTRs), establishing NUMTs as a novel source of VNTRs. In summary, the integrated analysis enhances our understanding of NUMT genomic architecture, population dynamics, and evolutionary implications in human and non-human primates, establishing NUMTs as dynamic genomic components of biomedical relevance.

## INTRODUCTION

Nuclear-embedded mitochondrial DNA segments (NUMTs) represent a unique class of genomic elements that chronicle the continuous transposition of mitochondrial DNA to the nuclear genome throughout evolutionary history^1–3^. Since the endosymbiosis event that established mitochondria, making mitochondrial DNA (mtDNA) the most proximal “foreign DNA” to the nucleus, mtDNA has been continuously integrated to the nuclear genome, potentially through intracellular processes involving mitochondrial degradation and DNA repair mechanisms^4^. Through evolution, some of these inserted mtDNA sequences have become permanent components of the nuclear genome, expressing mitochondrial genes using nuclear and cytoplasmic machinery and reinforcing the mitochondrion-nucleus codependency^2,4^. NUMTs, representing the other inserted mtDNA sequences that had been widely recognized as non-functional, are associated with more and more implications for genetics and disease as indicated in recent studies^5–7^.

A comprehensive inspection of NUMTs in large population datasets offers valuable resources for improving the understanding of their origin, evolution, genomic impact, and disease-related mechanisms. Previously, using next-generation sequencing data from the 100,000 Genomes Project in England, a study analyzed NUMTs in 66,083 individuals, revealing the ongoing NUMT formation^8^. Additionally, the study performed Oxford Nanopore sequencing for 39 individuals, showcasing the potential of long-read sequencing for NUMT characterization. However, the whole-genome landscape of NUMTs, particularly in complex genomic regions, has remained understudied^8–11^. This incomplete characterization has left critical knowledge gaps in NUMT genomic location, sequence features, and population frequency patterns, as well as the functional consequences of polymorphic NUMTs that vary among individuals. Likewise, the relationship between NUMT methylation levels and allele frequencies in human populations has not been comprehensively characterized, hindering our understanding of the methylation dynamics following NUMT insertion.

Recent advances in long-read sequencing and pangenome construction provide unprecedented opportunities to bridge the gaps in NUMT biology^12,13^. High-quality, haplotype-resolved genome assemblies from diverse populations enable precise identification of NUMTs in previously inaccessible genomic regions or those with complex concatenated structures, while pangenome graphs facilitate a comprehensive analysis of NUMT population genetics and evolutionary dynamics^14^. This technological revolution allows accurate distinction between fixed NUMTs present in all individuals and polymorphic variants exhibiting population-specific patterns, resolving the conflation inherent in single linear reference analyses. Furthermore, enhanced orthology alignments allow for precise NUMT characterization even within complex genomic regions^15,16^. This high-resolution characterization facilitates downstream functional analyses, such as genome-wide association studies (GWAS) and expression quantitative trait locus (eQTL) studies^17,18^. Additionally, by integrating pangenome-based NUMT characterization with comparative genomics across primate species, we can reconstruct the evolutionary history of NUMT insertion events and quantify lineage-specific insertion rates, thereby providing insights into cellular, reproductive, and developmental mechanisms driving NUMT insertion^19,20,21^.

Here, we employed a pangenome graph-based approach for comprehensive NUMT identification and applied it to a Minigraph-Cactus (MC) pangenome graph constructed from 538 haplotype-resolved assemblies across three consortia [269 individuals, including 160 from the Asian Pan-Genome Project Phase 1 (APGp1), 44 from the Human Pangenome Reference Consortium Phase 1^12^ (HPRCp1), and 65 from the Human Genome Structural Variation Consortium Phase 3^22^ (HGSVCp3)]. Additionally, we supplemented the pangenome analysis with short-read genotyping data from 2,504 unrelated individuals in the 1000 Genomes Project^23,24^ (1KGP). Our integrated analysis aims to: (1) generate a comprehensive map for human NUMTs, (2) characterize genomic features of fixed and polymorphic NUMTs, including mtDNA origin bias, localization preference, sequence divergence, and methylation dynamics, (3) explore functional impacts of polymorphic NUMTs on gene sequence, expression and splicing, and (4) delineate evolutionary dynamics of NUMTs across primates to understand the forces shaping NUMT insertion, retention, and expansion.

## RESULTS

### NUMT characterization in 269 human genomes

Conventional NUMT detection relies on short-read alignments to identify split reads exhibiting nuclear-mitochondrial dual-alignment clipping, followed by breakpoint refinement through realignment^8,25,26^ (Figure 1a). To overcome the inherent resolution and sensitivity limitations of the short-read approach, we developed a Pangenome Graph-based NUMT Detection (PG-NUMT) approach to comprehensively identify NUMTs across 200 + 1 benchmark individuals, including 40 from HPRCp1, 160 from APGp1, and the reference genome T2T-CHM13 (Figure 1b, Supplementary Table 1, Supplementary Figure 1-2, Methods). Briefly, PG-NUMT aligns multiple primate mitochondrial genomes to the pangenome graph, enabling sensitive, comprehensive NUMT detection with precise breakpoint identification. Using PG-NUMT, we identified a total of 1,122 NUMTs, including 918 derived from T2T-CHM13 (“reference NUMTs”) and 204 from the other genomes (“non-reference NUMTs”). In comparison, only 83 non-reference NUMTs were detected using the short-read approach, including two misidentified ones (Supplementary Table 2, Supplementary Figure 3). Benchmark results across the 200 individuals revealed a 2.52-fold improvement in NUMT detection sensitivity, with a particularly notable improvement for long NUMTs exceeding 1,000 bp (Figure 1c, 1d, from 5 to 19). Importantly, PG-NUMT can evaluate reference NUMT presence in any given individual and distinguish NUMTs between haplotypes, which is unattainable with short-read approaches (Figure 1a, 1b). Our benchmark results demonstrate that PG-NUMT substantially enhances both the accuracy and sensitivity of NUMT detection (Supplementary Table 2, Supplementary Figure 4).

**Figure 1.**
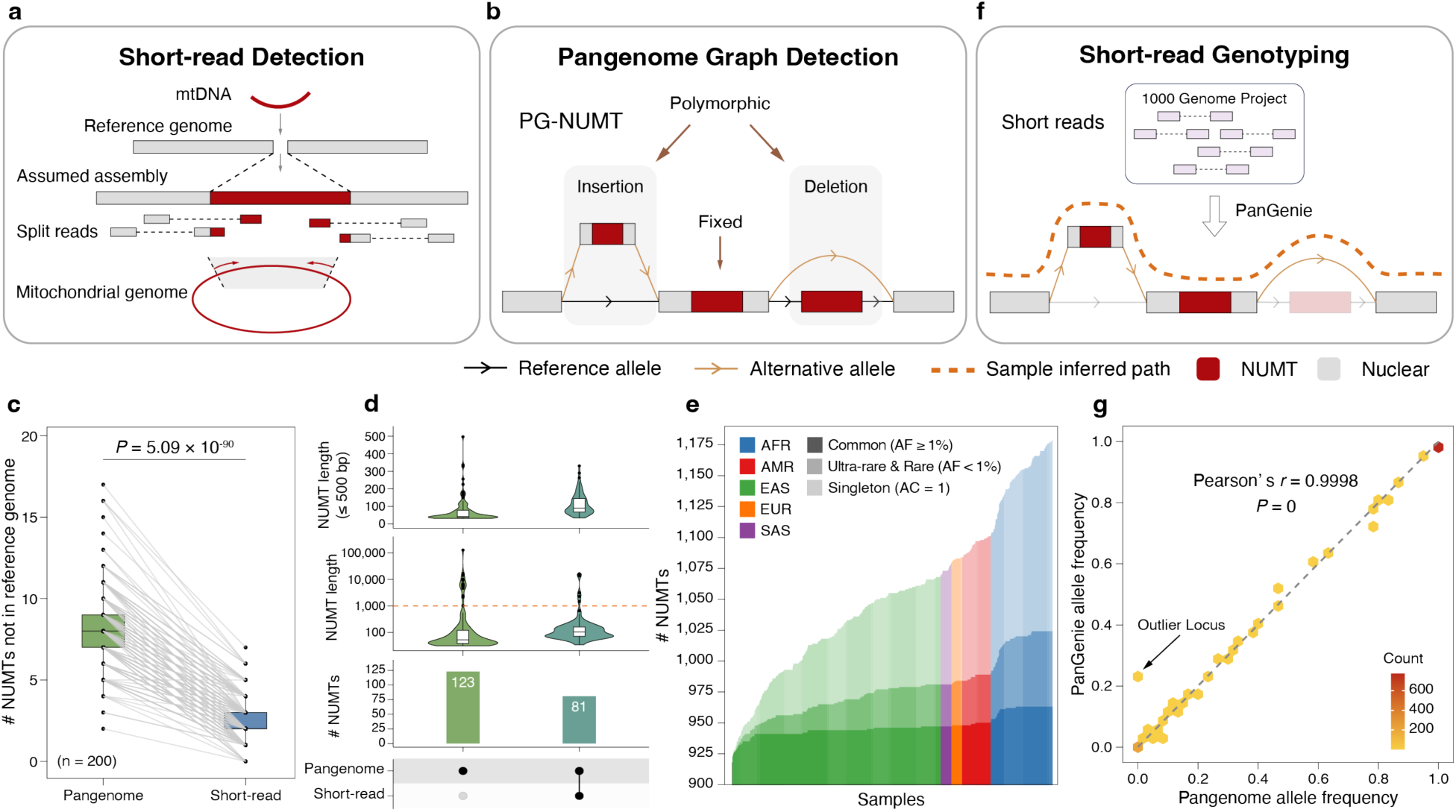
NUMT detection with pangenome graph. **(a)** NUMT detection using short-read sequencing data. **(b)** NUMT detection using the pangenome graph. PG-NUMT: Pangenome Graph-based NUMT Detection. **(c)** The paired comparison of detected NUMTs absent in the T2T-CHM13 reference genome shows a 2.52-fold increased sensitivity of PG-NUMT compared to the short-read approach (two-tailed Welch’s *t*-test, *P* = 5.09 × 10^-90^). Connecting lines indicate the same samples. **(d)** Distribution comparison of length and counts of identified NUMTs between PG-NUMT and the short-read approach. Left, NUMTs uniquely identified by PG-NUMT; Right, consensus NUMTs detected by both approaches, excluding two loci uniquely misclassified by short-read (Supplementary Figure 3). From bottom to top: UpSet plot comparing NUMT counts between approaches; violin plot of NUMT length distribution; violin plot of NUMT length distribution (≤ 500 bp). **(e)** Cumulative NUMT growth curves depicting sequential assembly addition to the pangenome graph, where colors from light to deep indicate singleton (AC = 1), ultra-rare & rare (AF < 1%), and common (AF ≥ 1%) NUMTs. **(f)** Short-read NUMT genotyping across 2,504 unrelated 1KGP individuals by PanGenie within a pangenome graph. **(g)** Comparison of allele frequencies between PanGenie genotypes and the pangenome graph across 40 HPRCp1 and 160 APGp1 individuals shows high concordance (Pearson’s *r* = 0.9998, *P* = 0). The outlier locus (pannumt_109, indicated by an arrow) exhibits reduced genotyping accuracy due to adjacent repetitive sequences and the inversion structure (Supplementary Figure 10).

Given its superior performance in benchmark evaluations, PG-NUMT was applied to a Minigraph-Cactus (MC) pangenome graph based on T2T-CHM13, which incorporates 269 individuals representing diverse global superpopulations (538 haplotype-resolved assemblies; Supplementary Table 1). Within the MC pangenome graph, 1,179 NUMTs were identified, including 918 reference and 261 non-reference NUMTs (Supplementary Table 3). Among the non-reference NUMTs, 7 were categorized as concatenated mega-NUMTs (NUMTs composed of multiple concatenated mtDNA-derived segments; Supplementary Table 3, Supplementary Figure 4-7). Each individual possesses an average of 8.67 non-reference NUMTs (s.d. = 2.85, Supplementary Figure 8). Among 1,056 NUMTs (excluding 123 in centromeric regions), 155 (14.7%) are singletons [allele count (AC) = 1], 61 (5.8%) are rare or ultra-rare NUMTs [AC > 1 and allele frequency (AF) < 1%], and 840 (79.5%) are common NUMTs (AF ≥ 1%), with all reference NUMTs classified as common. Notably, incorporating additional samples provides a marginal contribution to the pool of common NUMTs (Figure 1e), indicating that the current MC pangenome graph already captures a substantial portion of the common NUMT landscape in humans.

Our large, diverse pangenome graph improves NUMT detection resolution, particularly for common NUMTs that are extensively represented. This enables accurate NUMT genotyping through short-read sequencing in even larger human cohorts, thereby enabling further inspection of the NUMT landscape. Here, we genotyped NUMTs in short-read WGS data from 2,504 unrelated individuals within 1KGP using PanGenie^17^ based on the MC pangenome graph, successfully characterizing 1,043 loci (Figure 1f, Supplementary Table 3, Methods). To evaluate the genotyping performance, allele frequencies from the MC pangenome graph were compared with those generated by PanGenie (Methods). The high correlation between approaches confirms PanGenie’s reliability for NUMT genotyping accuracy across the 2,504 individuals (Pearson’s *r* = 0.9998, Figure 1g, Supplementary Figure 9-10).

### The fixed and population-stratified NUMTs in the human genome

Accurate and comprehensive NUMT characterization enhances the investigation of their population features. Here, NUMTs are classified into three categories based on allele frequencies and genome coordinates. Fixed NUMTs are stably integrated into the genomes with near-universal prevalence across populations (AF ≥ 95%), while polymorphic NUMTs exhibit individual-level variation (AF < 95%). Pericentromeric NUMTs, specifically located within centromeric regions, are classified as a separate category due to challenges preventing unambiguous classification as either fixed or polymorphic, such as suboptimal assembly accuracy and divergent alignments across genomes. Using PG-NUMT and PanGenie, we characterized 774 fixed NUMTs, 280 polymorphic NUMTs, and 123 pericentromeric NUMTs across 269 haplotype-resolved individuals and 2,504 unrelated short-read individuals (Figure 2a, Supplementary Table 3). All 774 fixed NUMTs are present in the reference genome T2T-CHM13. Among polymorphic NUMTs, 71.1% exhibit low allele frequencies in human populations [47.8% ultra-rare (AF < 0.1%) and 23.3% rare (0.1% ≤ AF < 1%)]. Notably, among the remaining 28.9% of polymorphic NUMTs [common NUMTs (AF ≥ 1%)], some exhibit population-specific allele frequency variations (Supplementary Table 4), hinting at different demographic histories or genetic drift across human populations (Methods).

**Figure 2.**
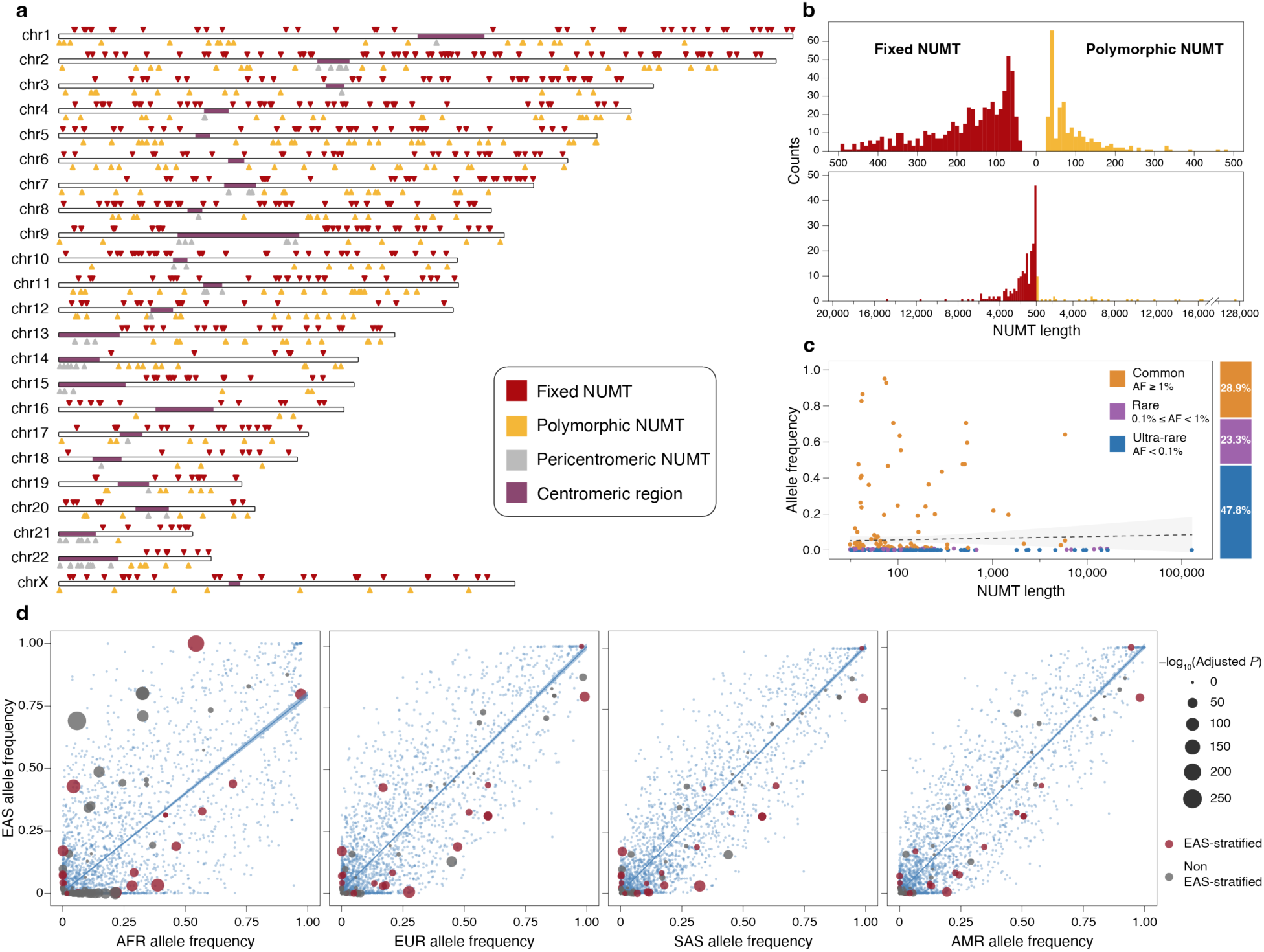
Overview of NUMTs in humans. **(a)** Ideogram of NUMTs from 578 human genome assemblies in this study. 774 fixed NUMTs are depicted above the ideogram track (red), while 280 polymorphic NUMTs (orange) and 123 pericentromeric NUMTs (grey) are shown below the ideogram track. Centromeric regions are labeled in purple. **(b)** Length distribution of fixed (left, red) and polymorphic (right, orange) NUMTs (top, 0-500 bp, 10-bp bin; bottom, > 500 bp, 200-bp bin). **(c)** The correlation between NUMT allele frequency and length is shown on the left. The proportions of NUMTs with different allele frequencies (ultra-rare (AF < 0.1%, blue), rare (0.1% ≤ AF < 1%, purple), and common (AF ≥ 1%, orange)) are shown on the right. **(d)** Allele frequency comparisons of NUMTs between East Asian (EAS) and other superpopulations. Circles represent NUMTs, with those colored red indicating EAS-stratified NUMTs (significantly different between EAS and other superpopulations at an adjusted *P* value < 0.05). Blue dots denote SNV allele frequencies across superpopulations, serving as controls.

Fixed NUMTs exhibit a length range of 38 bp to 14,855 bp (median: 203 bp; mean: 637.8 bp, Supplementary Figure 11), whereas polymorphic NUMTs range from 31 bp to an extraordinary 127,749 bp (median: 74.5 bp; mean: 1143.6 bp for all and 689.8 bp excluding the largest NUMT, Figure 2b). Despite this broad range, the majority of both fixed (73.4%) and polymorphic (87.9%) NUMTs are shorter than 500 bp. Unexpectedly, no significant negative correlation was observed between NUMT length and allele frequency, challenging previously reported patterns^8^ (Pearson’s *r* = 0.04, 95% CI: -0.08-0.15, *P* = 0.56, Figure 2c).

To further explore the population stratification patterns of NUMTs, we compared their distribution across human superpopulations among 2,504 unrelated 1KGP individuals. A total of 74 superpopulation-stratified NUMTs were identified (Figure 2d, Supplementary Table 4, Supplementary Figure 12, Methods). These population-stratified NUMTs likely represent independent genomic integration events during the evolutionary divergence of human populations.

### Genomic landscape of the fixed and polymorphic NUMTs

NUMT polymorphisms and allele frequencies may reflect the complex interplay of genetic drift, population dynamics and potential functional mechanisms that govern the evolutionary fate of different NUMTs. Integrating this perspective in NUMT analyses by characterizing fixed and polymorphic NUMTs separately helps reveal distinct genomic features, generating novel insights into the genomic effects of NUMTs. For instance, a remarkable bias was identified in the mtDNA origin distribution patterns. While polymorphic NUMTs exhibit uniform coverage across the mtDNA sequence (permutation test, *P* = 0.54, Methods), fixed NUMTs display a notably decreased coverage in the 3’-end region (chrM:72-573, mtDL3 hereafter, relative coverage < 0.30, permutation test, *P* = 0) of the non-coding displacement loop (D-loop) (Figure 3a, Supplementary Figure 13). To exclude the alignment artifacts as a potential cause for this observation, rigorous realignment analyses were performed using multiple mtDNA queries (N = 1,000)^27^, or reference sequences with shifted start positions (Supplementary Figure 14, Methods). Additionally, the divergent patterns observed between fixed and polymorphic NUMTs suggest that the reduced coverage in mtDL3 more likely arises from post-insertion events rather than bias during integration (Figure 3a). To explore the mechanism underlying the selective removal, we assessed the regulatory potential of the mtDL3-derived sequence through a reporter assay (Figure 3b-3d, Supplementary Figure 14, Methods). The mtDL3-derived sequence exhibits weak *cis*-regulatory activity, while neither the negative control nor the mtDL5-derived sequence (the 5’-end region of the D-loop, chrM:16,024-16,569) exhibits such activity (Figure 3c, 3d). The potential modulatory effect of mtDL3-derived sequences provides a potential biological basis for their preferential elimination from fixed NUMT populations. However, the extent to which this activity drives their selective removal remains unclear and requires additional investigation.

**Figure 3.**
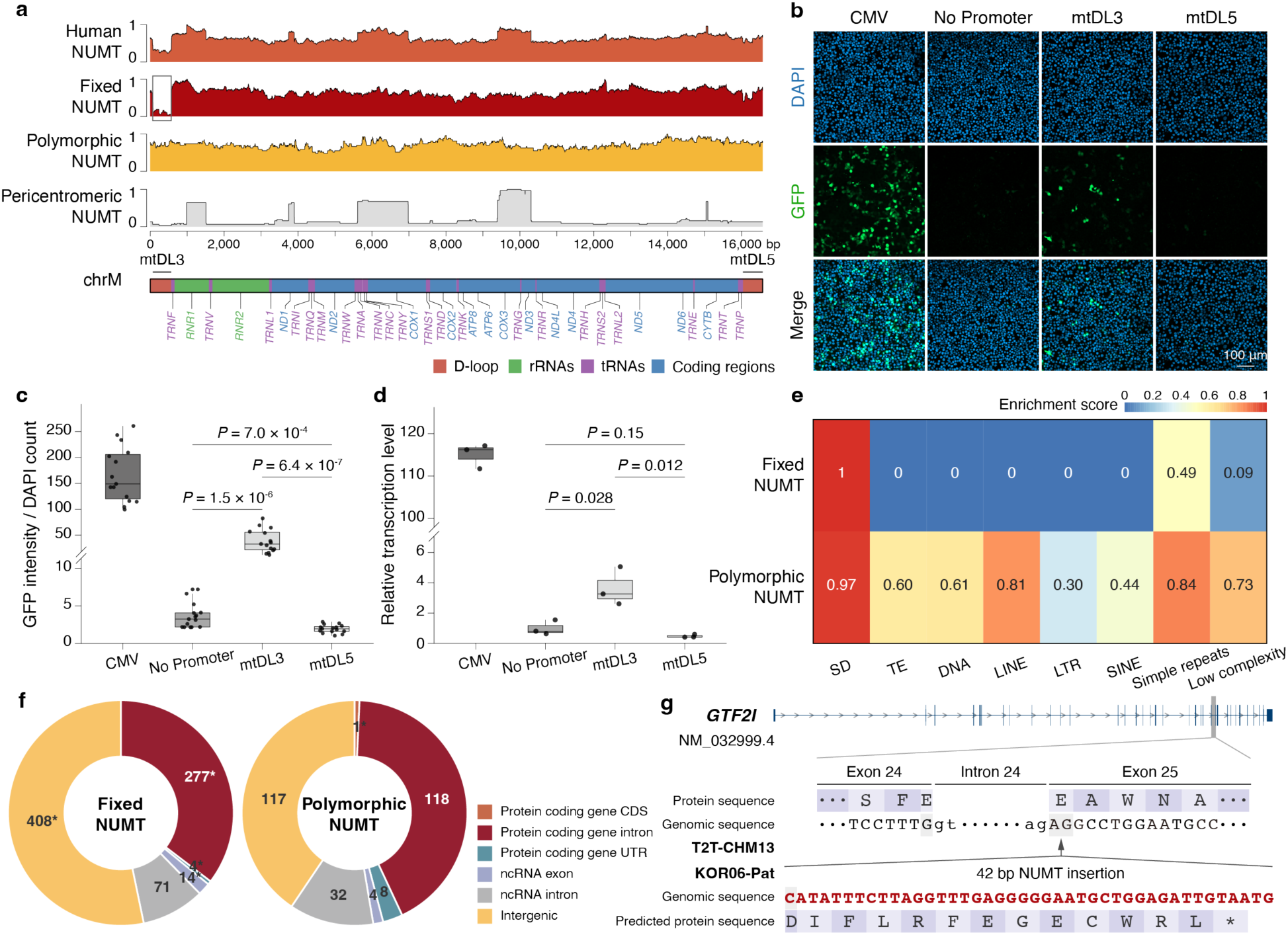
Distributions of fixed and polymorphic NUMTs. **(a)** NUMT coverage relative to the mtDNA coordinates. The x-axis represents the nucleotide positions of mtDNA, while the y-axis indicates the relative coverage distribution across distinct NUMT categories (normalized by their maximum coverages). From top to bottom: all NUMTs, fixed NUMTs, polymorphic NUMTs, and pericentromeric NUMTs. The mitochondrial gene annotation is presented at the bottom. The mtDL3 region (chrM:72-573) and the mtDL5 region (chrM:16,024-16,569) are marked with grey lines (Supplementary Figure 14). Reduced coverage in the mtDL3 region of fixed NUMTs is highlighted by the grey box. **(b)** Representative fluorescence images of HEK293T cells transfected with recombinant plasmids, illustrating the in *vitro* promoter activity of mtDL3 and mtDL5. Images in the CMV group were acquired using substantially lower laser excitation intensity to avoid overexposure. DAPI, 4’,6-diamidino-2-phenylindole; GFP, green fluorescent protein; CMV, cytomegalovirus enhancer and promoter. **(c)** Quantitative analysis of the fluorescence signals shown in Figure 3b. Five images were acquired for each of three independent transfection replicates. **(d)** Relative transcriptional activity of mtDL3 and mtDL5, measured by eGFP expression. Three qPCR technical replicates were performed for each of three independent transfection replicates. **(e)** Heatmap of enrichment analysis of fixed (top) and polymorphic (below) NUMT insertions across repetitive elements in the human genome. Values represent enrichment scores derived from permutation tests relative to random expectations (Methods). SD, segmental duplication; TE, transposable element; DNA, DNA transposon; LINE, long interspersed nuclear element; LTR, long terminal repeat; SINE, short interspersed nuclear element. **(f)** Pie chart illustrating the proportion of fixed (left) and polymorphic (right) NUMT insertions across different genomic regions. CDS, coding sequence; UTR, untranslated region. The asterisk (*) indicates a significant deviation from the null distribution. **(g)** A NUMT insertion potentially alters the CDS of *GTF2I*. The gene model of *GTF2I* (NM_032999.4) is presented at the top, with the annotation from T2T-CHM13 (reference). The 42-bp NUMT insertion sequence within the KOR06-Pat (NUMT insertion carrier) genome is highlighted in red, an in-frame insertion predicted to cause premature termination. The asterisk (*) indicates the predicted termination codon.

Additionally, the mtDNA origin analysis also reveals pericentromeric NUMT enrichment in five specific mtDNA regions (relative coverage > 0.50, chrM:986-1,509; chrM:3,744-3,897; chrM:5,612-6,979; chrM:9,393-10,302; chrM:15,037-15,087, Figure 3a, Supplementary Figure 9). Since 81.43% of the length of these regions overlaps with segmental duplications, their elevated coverage is potentially attributable to genomic duplication events.

Next, we analyzed the localization preference of fixed and polymorphic NUMTs in the human genome. Enrichment analysis reveals that both fixed and polymorphic NUMTs are significantly enriched within or near segmental duplications (fixed NUMTs, permutation test, *P* = 0; polymorphic NUMTs, permutation test, *P* = 0.03; Figure 3e, Methods). Fixed NUMTs are rarely found in regions near or within transposable elements, including DNA transposons, LINEs, LTRs, and SINEs (permutation test, *P* = 0, Supplementary Figure 15). In contrast, polymorphic NUMTs display no significant deviation from the null distribution in association with transposable elements (permutation test, *P* = 0.40). These findings reveal distinct localization preferences between fixed and polymorphic NUMTs, implicating roles of duplications and transposable elements in shaping their genomic distribution (Figure 3e, Supplementary Figure 16-17).

The integration of NUMTs into the nuclear genome may disrupt gene structure and regulation, potentially affecting protein function^5,6^. To explore potential function-related effects, the distribution of fixed and polymorphic NUMT insertions was examined across different genomic regions. Fixed NUMTs showed significant enrichment in intergenic regions (permutation test, *P* = 0, Figure 3f), representing the genomically “safe” locations where insertion is least likely to disrupt essential cellular functions. In contrast, polymorphic NUMTs showed no significant location bias (permutation test, *P* = 0.50), indicating that such location bias is established through post-integration selective mechanisms over evolutionary time rather than insertion location preference (Figure 3f, Supplementary Figure 18). Notably, one singleton 42-bp NUMT inserted into the coding sequence (CDS) of *GTF2I* was identified among 280 polymorphic NUMTs (Figure 3f, 3g). *GTF2I* encodes a transcriptional factor and is located within a deletion dominantly associated with Williams-Beuren syndrome^28^. The insertion occurs after the first nucleotide in exon 25 but is not predicted to affect splicing by SpliceAI^29^ (delta scores for acceptor loss = 0.07; delta scores for acceptor gain, donor gain, and donor loss = 0.00). Nanopore long-read RNA-seq reads with the NUMT insertion, coupled with the absence of exon-skipped reads, confirm an in-frame insertion leading to premature termination and nonsense-mediated decay (Figure 3g, Supplementary Figure 19). However, considering the carrier’s apparent lack of disease phenotype, further studies are required to investigate any potential physiological consequences or compensatory mechanisms.

### Sequence features and methylation patterns of NUMTs

The nuclear environment imposes fundamentally different mutational constraints compared to the mitochondrial genome, leading to progressive sequence divergence that serves as a molecular clock for estimating NUMT integration timing^8^. We found that fixed NUMTs exhibit significantly lower sequence identity with mtDNA compared to polymorphic NUMTs (two-tailed Welch’s *t*-test, *P* = 7.67 × 10^-213^, Figure 4a, Supplementary Figure 20), consistent with the expectation that fixed NUMTs have accumulated more mutations during their prolonged residence within the nuclear genome.

**Figure 4.**
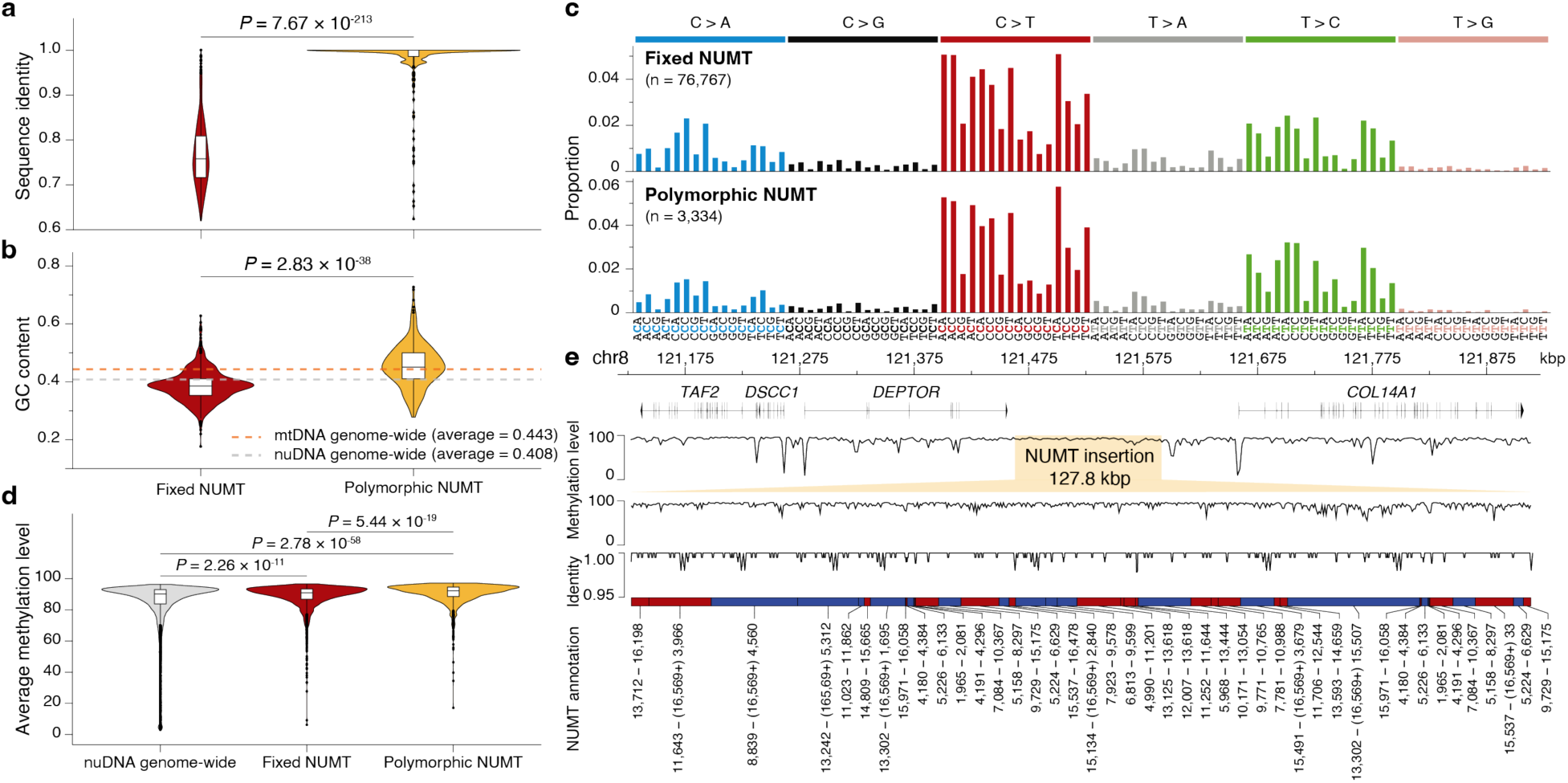
Sequence characteristics of fixed and polymorphic NUMTs. **(a)** Comparison of sequence identity to mtDNA between fixed and polymorphic NUMTs. **(b)** Comparison of GC content between fixed and polymorphic NUMTs. The orange and grey lines represent the GC content observations of random sequences at the mitochondrial and nuclear genome-wide levels, respectively. **(c)** Trinucleotide mutation spectrum of fixed (top) and polymorphic (bottom) NUMTs compared with the revised Cambridge Reference Sequence (rCRS). **(d)** Average methylation level distribution of NUMT windows among fixed NUMTs (red), polymorphic NUMTs (yellow), and genome-wide nuclear DNA (grey) (two-tailed Mann–Whitney *U*-test). Each NUMT was divided into 200-bp windows, and only NUMTs greater than 1 kbp were analyzed. **(e)** Methylation pattern of a mega-NUMT on C060-CHA-N20#Mat. The top panel displays methylation levels across the mega-NUMT (chr8:121,463,091-121,590,920) and the flanking regions (chr8:121,126,410-121,914,383). The middle panel displays sequence identity to mtDNA. The bottom panel illustrates the concatenated NUMT structure with corresponding mitochondrial coordinates, where red and blue regions denote forward and reverse mtDNA strands, respectively. Coordinates formatted as "(16,569+) X" represent regions spanning the nucleotide position “1” of the mtDNA reference.

In addition, fixed NUMTs exhibit significantly reduced GC content compared to both polymorphic NUMTs (two-tailed Welch’s *t*-test, *P* = 2.83 × 10^-38^, Figure 4b) and the mitochondrial genome-wide average (permutation test, *P* = 0), with a GC content of approximately 5.38% lower than the nuclear genome-wide average (from 0.408 to 0.386). This pattern aligns with mutation spectrum analyses, which reveal a pronounced preferential bias of C→T mutation in both fixed and polymorphic NUMTs (Figure 4c). However, the mutation spectra do not differ significantly between fixed and polymorphic NUMTs (Supplementary Table 5), suggesting shared post-integration mutational processes in both categories consistent with nuclear CpG transitions^30^. These observations imply that the observed differences in sequence identity and GC content between NUMT categories primarily reflect the cumulative effects of time rather than distinct mutational mechanisms.

The human nuclear genome is typically hypermethylated in most inactive genomic regions^31,32^, while the entire mitochondrial genome exhibits low-to-none methylation levels^33,34^. Considering this distinction, methylation analyses for NUMTs may reveal properties related to their insertion and post-insertion mechanisms. Contrasting sharply with mtDNA, both fixed and polymorphic NUMTs exhibit high methylation levels across most genomic windows (200-bp windows; fixed NUMTs: mean = 88.41%, median = 90.89%; polymorphic NUMTs: mean = 90.46%, median = 92.23%, Figure 4d, Methods). Considering methylation levels are comparable between fixed and polymorphic NUMTs despite their presumably different evolutionary ages, the related epigenetic response is likely to be both rapid and stable, thereby effectively masking age-dependent differences. Moreover, uniform patterns of methylation level and sequence identity persist across all constituent segments of a concatenated mega-NUMT (Figure 4e). These observations align with the proposed formation mechanism in which multiple mtDNA fragments are simultaneously integrated into the nuclear genome during NUMT insertion^25^.

### Assessment of modulatory effects of polymorphic NUMTs

The potential for polymorphic NUMTs to influence gene function represents a critical frontier in understanding how mtDNA insertions contribute to human phenotypic diversity and disease susceptibility. Unlike fixed NUMTs, which have undergone extensive evolutionary selection, polymorphic NUMTs may retain modulatory potentials due to their recent integration and variable presence across populations.

To investigate the potential gene function modulation roles of polymorphic NUMTs, we performed *cis*-eQTL and *cis*-sQTL analysis of polymorphic NUMTs, as well as SNVs in their flanking 500-kb regions using RNA-seq data from 731 individuals in the MAGE dataset^35^. We identified 7 significant eNUMT-eGene and 16 sNUMT-sGene pairs, where eNUMTs/sNUMTs and eGenes/sGenes represent NUMTs contributing to *cis*-eQTL/sQTL signals and their corresponding target genes (Supplementary Table 6-7). Among those, three NUMTs exhibited dual associations impacting both gene expression and splicing, including pannumt_14 (associated with *GNL2*, Figure 5a-5c), numt_353 (associated with *RASGRP3*, Figure 5d-5f), and numt_310 (associated with *SMAD2*, Figure 5g-5i).

**Figure 5.**
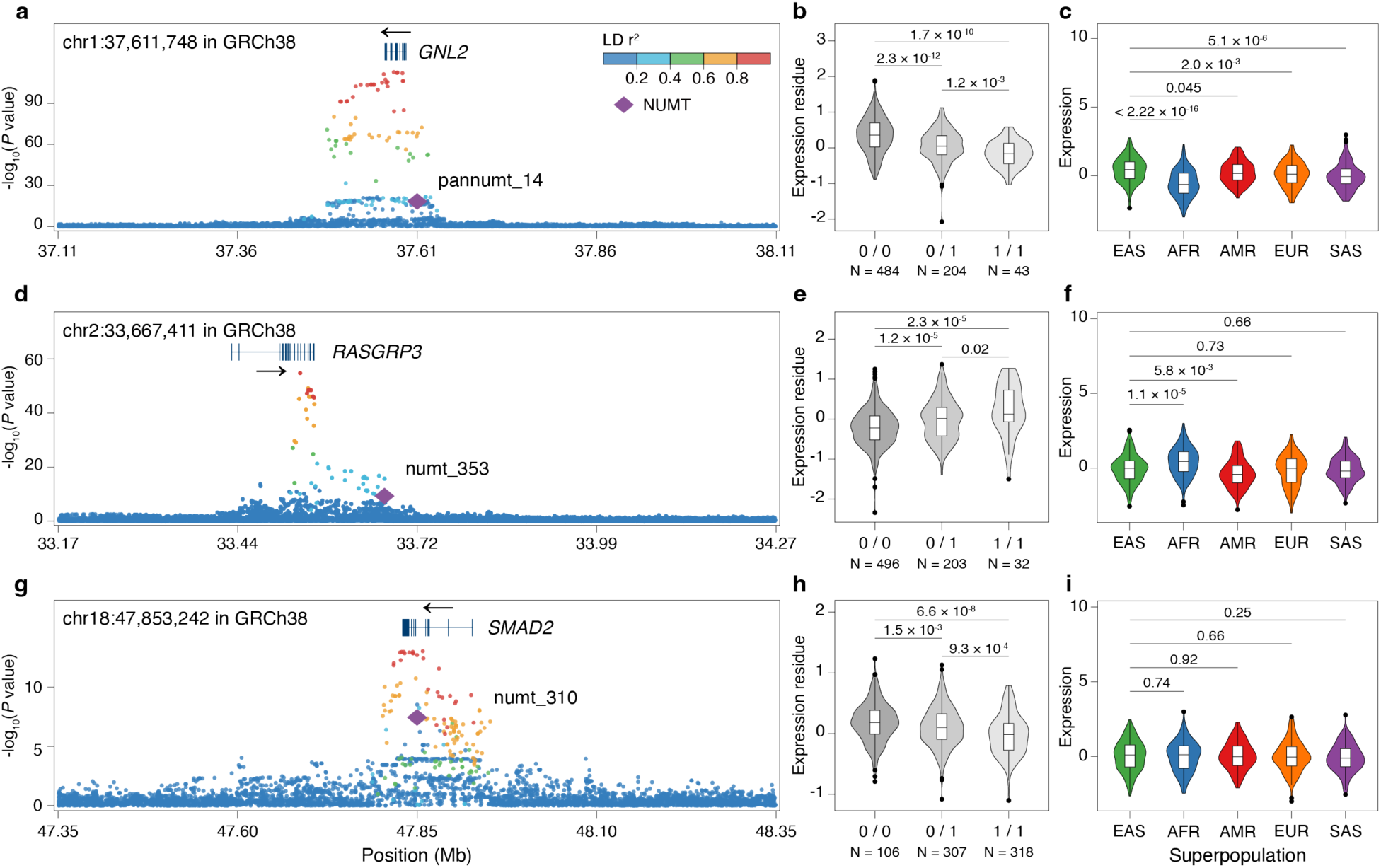
Significant *cis*-eQTL effects of NUMTs. **(a, d, g)** LocusZoom plots of regional eQTL association analyses for three NUMTs exhibiting dual QTL effects, including pannumt_14 and *GNL2* (a), numt_353 and *RASGRP3* (d), numt_310 and *SMAD2* (g). Arrows indicate the direction of gene transcription. **(b, e, h)** Covariate-adjusted normalized expression residuals comparisons across NUMT genotypes for *GNL2* (b), *RASGRP3* (e), and *SMAD2* (h). **(c, f, i)** Normalized expression comparisons across superpopulations for *GNL2* (c), *RASGRP3* (f), and *SMAD2* (i). Sample sizes of each superpopulation: EAS (N = 141), AFR (N = 196), AMR (N = 113), EUR (N = 142), and SAS (N = 139).

Specifically, pannumt_14 represents a 71-bp NUMT located in the 5’ upstream region of G Protein Nucleolar 2 (*GNL2*, a nucleolar GTP-binding protein required for nuclear export and maturation^36^) and is seen in conjunction with decreased expression (Figure 5a, 5b). Similarly, numt_310 also showed reduced expression association (Figure 5g, 5h), which represents a 210-bp NUMT within the intron of SMAD Family Member 2 (*SMAD2*), a gene encoding the signaling transducer and transcriptional factor that regulates cell proliferation, apoptosis, and differentiation through TGF-β signaling pathway^37^. By contrast, we identified an enhanced expression correlation between numt_353, a 245-bp NUMT insertion and its adjacent gene RAS Guanyl-Releasing Protein 3 (*RASGRP3*) (Figure 5d). *RASGRP3* encodes a guanine nucleotide exchange factor that activates the oncogenes *HRAS* and *RAP1A*, reported to be associated with systemic lupus erythematosus and several cancers^38^. Therefore, while none of these NUMTs constituted the lead QTL variants in their respective genomic loci (Figure 5a, 5d, 5g, Supplementary Figure 21), the crucial importance of these three genes warrants further functional examination of whether the presence/absence of these NUMTs possesses gene modulatory potential.

Interestingly, these three NUMTs also exhibit significantly lower allele frequencies in EAS than in other superpopulations (Supplementary Table 4), suggesting that these variants may reflect population-specific demographic histories or represent founder effects during human population expansion. However, population-specific expression differences corresponding to NUMT frequency patterns were observed only for *GNL2* (Figure 5c) and *RASGRP3* (Figure 5f), but not for *SMAD2* (Figure 5i), indicating that the interplay of NUMT presence, population stratification, and functional consequences is not straightforward and likely depends on additional genetic and environmental factors.

### NUMT evolution in primates

Although previous studies revealed the continuous integration of NUMTs during primate evolution, the lack of high-quality genomic resources has hindered a deeper understanding of NUMT phylogenetic patterns and insertion rates across primate lineages^11,20,39^. Here, we utilized 20 high-quality long-read non-human primate genomes (together with 270 human genomes) to systematically identify NUMTs, classify them as either shared or lineage-specific, and estimate the NUMT insertion rate in evolutionarily conserved syntenic regions for each lineage (Supplementary Table 8-9, Figure 6a, Supplementary Figure 22). NUMT insertion events exhibit a significant positive correlation with the evolutionary genetic distances between species, as measured by the SNV rate (Pearson’s *r* = 0.83, *P* = 2.71 × 10^-3^, Supplementary Figure 23).

**Figure 6.**
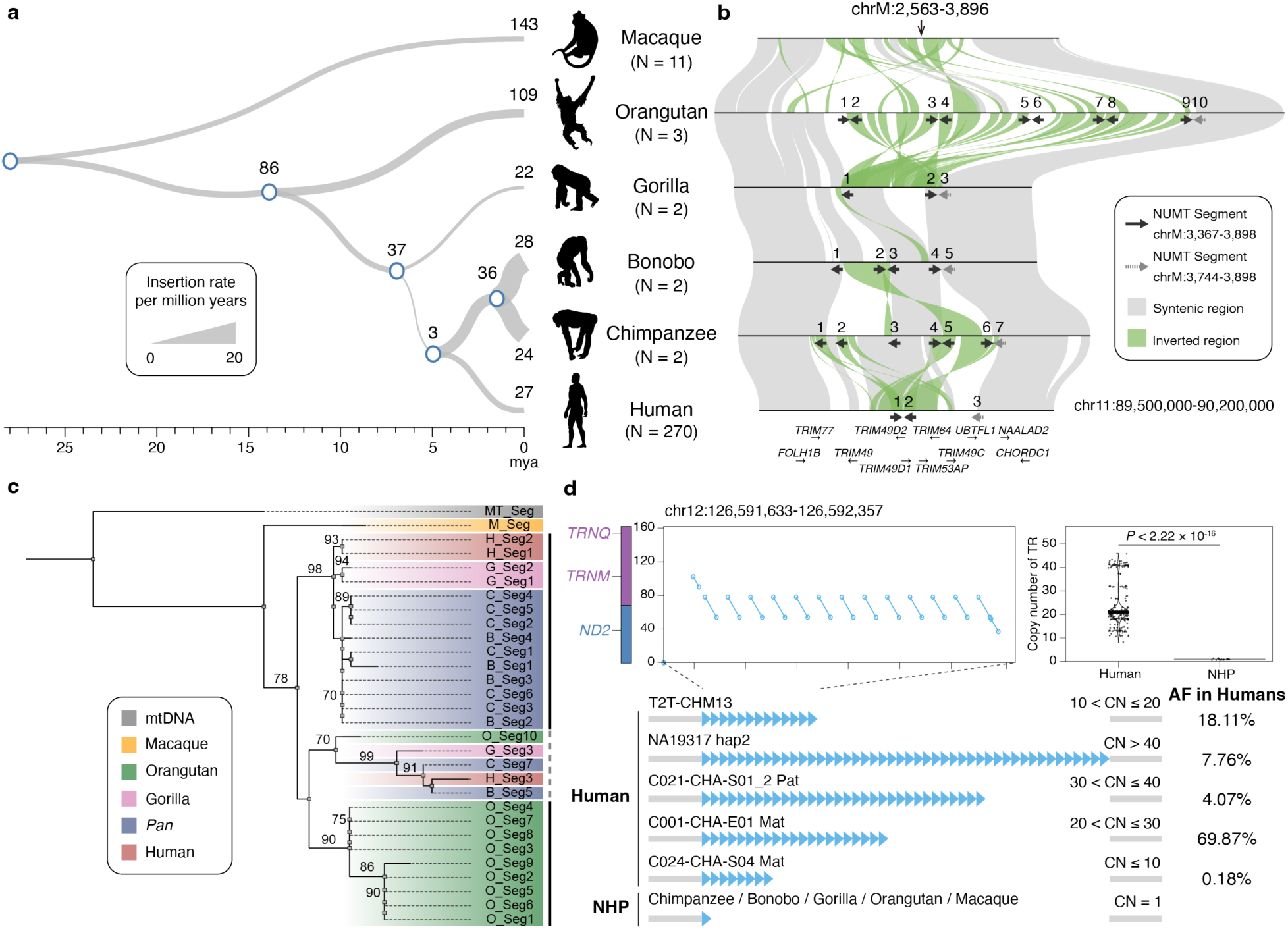
NUMT evolution in primates and VNTR origins within NUMTs. **(a)** The phylogenetic tree of fixed lineage-specific NUMTs in syntenic regions in primate evolution. Tree branch widths are scaled proportionally to the NUMT insertion rate per million years for each lineage (ranging from 1.5 to 18.7 insertions per million years). The number of NUMT insertions for each lineage is shown at each node or tip. **(b)** Syntenic comparison of recurrent NUMTs across primates at chr11:89,500,000-90,200,000 in T2T-CHM13. Black and grey arrows indicate NUMT segments corresponding to human chrM:3,367-3,898 and chrM:3,744-3,898, respectively. Arrow orientation reflects the direction of the NUMT segments. The gene models are present at the bottom. **(c)** The phylogenetic tree of NUMT segments from Figure 6b illustrates NUMT duplication and lineage-specific expansion. Bootstrap values (bootstrap value ≥ 70) are indicated at branch nodes. **(d)** A human-specific NUMT-derived VNTR. The syntenic comparison of the NUMT sequence and mtDNA sequence (left) and tandem repeat motif copy number distribution (right) are shown at the top. Schematic representations of the primate-specific NUMT-derived VNTR and its allele frequency distribution across humans are displayed at the bottom.

Furthermore, the observed NUMT accumulation pattern is consistent with lineage-specific insertion events^19^ (Supplementary Figure 23). Notably, the *Pan* lineage exhibits higher NUMT insertion rates in syntenic genomic regions across primates (chimpanzees, 16.0 insertions per million years; bonobos, 18.7), whereas gorillas display a comparatively lower insertion rate (3.1 insertions per million years). In addition to these syntenic regions, 8 recurrent duplication clusters were identified in evolutionarily dynamic regions across primates (Figure 6b, Supplementary Figure 24, Supplementary Table 10), with most prominent example located at chr11:89,828,225-90,018,356 in T2T-CHM13 (Figure 6b). Phylogenetic analysis of these NUMT segments across primates reveals that duplication events occurred independently in both the *Pan* and gorilla lineages (Figure 6c). Another distinct expansion pattern was observed in macaques, where *DUX4*-NUMT joint tandem duplication generated two macaque-specific arrays (chr1 and chr6, Supplementary Figure 25-26). This complex structure disrupts the continuous high-methylation region typically observed in humans. These findings indicate that duplication serves as a driving force behind NUMT expansion during primate evolution, a pattern similarly observed in human populations (Figure 3e, Supplementary Figure 16-17). This duplication-driven expansion model provides a mechanistic explanation for the clustered distribution of certain NUMTs observed in human and non-human primate genomes and suggests that NUMT abundance may be influenced by both post-integration duplications and initial rates of mitochondrial DNA transfer. Additionally, fixed NUMTs in non-human primates consistently exhibit decreased coverage in the mtDL3 region across the mtDNA sequence, mirroring the pattern observed in human fixed NUMTs (Supplementary Figure 27). This phylogenetic conservation of sequence bias suggests that the selective pressures responsible for mtDL3 depletion have operated throughout primate evolution, indicating that certain mitochondrial sequences possess intrinsic properties that make them less suitable for long-term nuclear residence.

### The origins of VNTR within NUMTs

Previous comparative genomic analyses reveal that homologous NUMT loci accumulate sequence variations throughout evolution, including SNVs and indels^39^. Additionally, our comparative analysis reveals a previously unrecognized source of genomic variation through the identification of NUMT-derived variable number tandem repeats (VNTRs), establishing mitochondrial insertions as contributors to the diversity of complex genomic regions. We identified two distinct NUMT-derived VNTRs with different evolutionary histories and potential functional significance, which are a human-specific NUMT-derived VNTR (Figure 6d) and a NUMT-derived VNTR expanded in humans (Supplementary Figure 28). The human-specific NUMT-derived VNTR (chr12:126,591,633-126,592,357 in T2T-CHM13) resides within a “gene desert” region (68,542 bp from the nearest gene *LOC100996671*). This VNTR displays a copy number variation ranging from 8 to 46 in humans and has expanded in archaic humans (including both the *Neanderthal* and the *Denisovan*; CN > 1; Supplementary Figure 29) but remains a single-copy locus in non-human primates. Notably, 2-bp “AT” microhomologies are observed at the repeat boundaries, suggesting that replication slippage might be associated to this expansion (Supplementary Figure 30). Similarly, the NUMT-derived VNTR at chr2:238,011,642-238,015,272 in T2T-CHM13 exhibits a copy number range of 9 to 18 in humans, representing an expansion compared to the fewer than 7 copies typically observed in non-human primates (Supplementary Figure 28). Interestingly, this VNTR resides within an intron of *MLPH* (encoding melanophilin), a gene potentially associated with human pigmentation^40^ and Griscelli Syndrome 3^41^. The discovery that mitochondrial sequences can serve as substrates for VNTR formation adds a new dimension to our understanding of how repetitive elements contribute to genome evolution and suggests that the functional impact of NUMT insertions may extend beyond their initial integration sites through subsequent structural rearrangements.

## DISCUSSION

This comprehensive investigation of nuclear-embedded mitochondrial DNA sequences represents the most extensive characterization of NUMT landscapes in human and primate genomes to date, fundamentally advancing our understanding of mitochondrial-nuclear genome interactions and their evolutionary consequences. By integrating population genomics, comparative phylogenetics, and functional genomics approaches, we have revealed that NUMTs constitute a dynamic and functionally relevant component of primate genome evolution, challenging views of these sequences as merely genomic fossils and establishing them as potential active contributors to genetic diversity, gene modulation, and evolutionary adaptation.

Long-read sequencing and pangenome graphs enable the precise identification of NUMT insertion sites and sequence characteristics. Moreover, this approach enhances comparative analyses of NUMT diversity by effectively distinguishing fixed NUMTs from polymorphic NUMTs in human populations. Fixed and polymorphic NUMTs exhibit both shared and divergent patterns in mtDNA origin bias, genomic insertion preference, sequence divergence, and methylation levels, which aid in studying post-integration modification processes and selective constraints on NUMT insertion. Notably, contrary to what was previously reported^8^, no significant negative correlation was observed between NUMT length and allele frequency in this study (Figure 2c). Therefore, it is likely that the selective removal of certain NUMTs is more associated with factors other than NUMT length, such as insertion position and insertion sequences. Additionally, high-quality genomes unlock NUMT detection within previously uncharacterized regions, including centromeric regions. In this study, it is revealed that pericentromeric NUMTs are enriched in human acrocentric chromosomes (chr13, chr14, chr15, chr21, and chr22, permutation test, *P* = 0), suggesting the enrichment in specific mtDNA regions likely originates from dynamic recombination events among both homologous and heterologous acrocentric chromosomes^42^. Additionally, high-quality genomes enable precise identification of 7 mega-NUMTs (maximum length 127.7 kbp), revealing their complex structures.

The conservation of sequence-specific selective pressures across primate evolution, particularly the consistent depletion of mtDL3-derived sequences in fixed NUMTs, reveals universal constraints on mitochondrial-nuclear genome compatibility that may have operated for millions of years (Figure 3a, Supplementary Figure 27). This phylogenetic conservation implies that certain molecular mechanisms responsible for NUMT sequence evolution might be fundamental to eukaryotic biology rather than species-specific adaptations. Understanding these constraints could provide additional insights into the cellular processes that govern foreign DNA integration and retention, with implications for both the understanding of natural genome evolution and the design of therapeutic genetic interventions. Interestingly, mtDL3 also exhibits low constraint within the mitochondrial genome^43^ (Supplementary Figure 31). This raises the question of whether the observed decreased mtDL3 coverage in fixed NUMTs may be an analytical artifact. Considering (1) mtDL3 coverage is not decreased in polymorphic NUMTs and (2) realignment using multiple mtDNA references^27^ confirms the decrease in fixed NUMTs, we hypothesize that the selected removal of mtDL3 in fixed NUMTs should have a biological cause (Supplementary Figure 14). However, whether the overlap of low coverage in fixed NUMTs and reduced constraint in the mitochondrial genome is coincidental or biological requires additional mechanistic studies. Moreover, the compromised stability of genomic DNA may also be associated with selective removal, warranting follow-up in future studies.

The precise characterization of NUMTs provides novel insights into the mechanisms underlying their insertion. For *de novo* NUMT insertions, previous studies have highlighted the critical role of double-strand break (DSB) repair mechanisms, particularly microhomology-mediated end joining (MMEJ) in *de novo* NUMT insertion^8^. In addition to that, our analyses provided substantial direct evidence revealing that segmental duplication of pre-integrated NUMTs serves as a key mechanism driving NUMT expansion across human populations and throughout primate evolution. These findings propose that NUMT dynamics may involve initial *de novo* insertions mediated by mitochondrial DNA transfer to the nucleus, followed by duplication events within nuclear genomes. This process model better elucidates the mechanisms governing NUMT formation and explains the polymorphic variations observed across populations and evolutionary lineages.

Considering the potential genomic impacts NUMT insertion may generate, it is of significant interest to identify the insertion locations and associated gene-related effects. Fixed NUMTs are stably integrated into genomes, predominantly localized in non-functional genomic regions. In contrast, polymorphic NUMTs show greater potential to exert functional consequences. Both prior research and our current analyses indicate that while polymorphic NUMTs may disrupt protein-coding genes and cause diseases^5,6^, these variants occur infrequently in human populations as a result of negative selection. Common polymorphic NUMTs display heterogeneous allele frequency distributions across populations. Notably, certain NUMT variants exhibit *cis*-eQTL and/or *cis*-sQTL effects and colocalize with lead SNPs in shared linkage disequilibrium (LD) blocks. Importantly, while fine-mapping analysis cannot directly indicate the function of NUMTs, it does not exclude their modulatory effects within these LD blocks. These potential modulatory roles require validation through precise experimental evidence.

The combination of high-quality long-read genomes^44–46^ with advanced whole-genome alignment tools^47^ has enhanced cross-species comparisons among primates, generating higher-resolution lineage-specific NUMT datasets than those reported in prior studies^11,20,39^. The observed divergence in lineage-specific NUMT insertion rates across species likely stems from different frequencies of mtDNA transfer events and lineage-specific selective pressures, which may be associated with interspecies variations in mitochondrial architecture and reproductive strategies^48^. Moreover, comparative analyses also revealed two NUMT-derived VNTRs, with divergent nuclear-mitochondrial genome constraints likely underlying their formation in these mtDNA-derived segments. Finally, considering cross-species NUMT comparisons offer critical insights into characteristics and mechanisms of NUMT integration, high-quality long-read genomes across the broader primate phylogeny are needed to achieve more comprehensive evolutionary analyses.

## Code availability

Custom scripts used in this study are available at GitHub (https://github.com/LiantingFu/NUMT_Analysis).

## Acknowledgments

We thank Nahyun Kong and Dr. Sheng Chih Jin for their suggestions for this study. We thank the HPRC and Primate T2T Consortium for providing the long-read human and great ape genome assemblies. The computations in this study were run on the Siyuan-1 supported by the Center for High Performance Computing at Shanghai Jiao Tong University.

## Author contributions

L.F., K.M. and Y.M. conceived the project; L.F. performed NUMT detection in humans and NUMT evolutionary analysis across primates; L.F., J.C., D.L., S.D., Z.W., H.M., Z.L., N.L., X.Y., Y.S. and K.M. contributed to NUMT genomic feature analysis; J.C., S.D. and Y.S. contributed to QTL analysis of polymorphic NUMTs; D.L., Z.W., H.M., Z.L. and K.M. contributed to the analysis of NUMT selective removal mechanisms; and D.W., C.Y. and G.Z. generated the genome assemblies of APG individuals. L.F., J.C., G.Z., K.M., and Y.M. drafted the manuscript. All authors read and approved the manuscript.

