## Supplemental Figures for "A long-read human pangenome initiative for comprehensive interpretation of nuclear-embedded mitochondrial DNA"

For

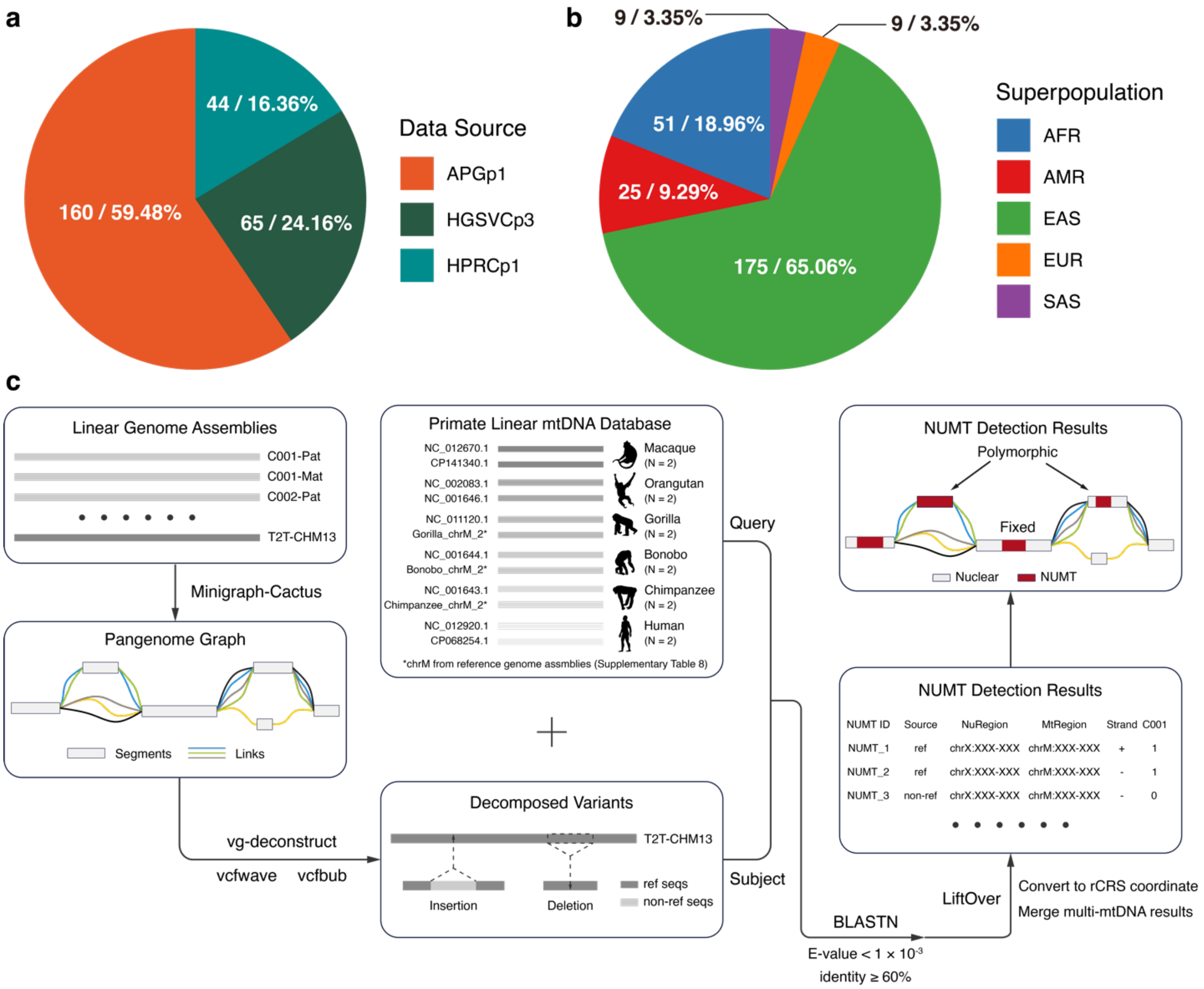


**Supplementary Figure 1.** The data source **(a)** and superpopulations **(b)** classifications of individuals in the study, respectively. The number of individuals and their proportions for each category are displayed in the pie chart. **(c)** The workflow of the PG-NUMT method.

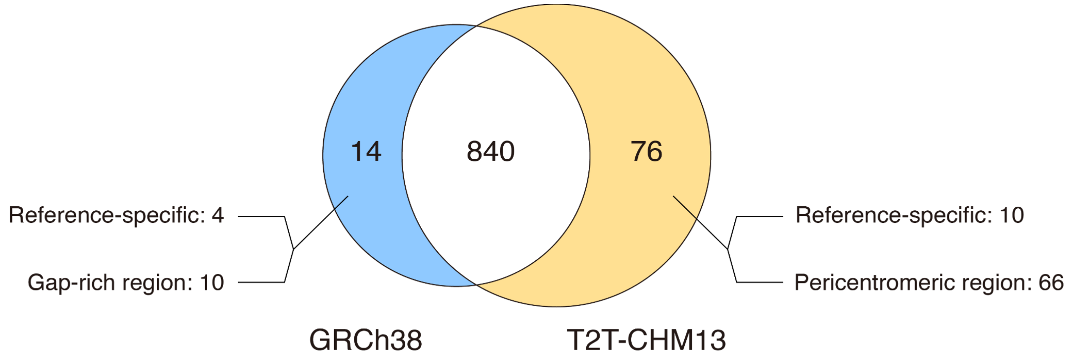


**Supplementary Figure 2.** NUMT comparison between GRCh38 and T2T-CHM13. While 840 NUMTs are conserved between assemblies, T2T-CHM13 contains 76 unique NUMTs, with 66 attributed to its superior resolution in pericentromeric regions and 10 to haplotype specificity. Conversely, GRCh38 contains 14 unique NUMTs, of which 4 are attributed to haplotype variation, and 10 are located in the gap-rich regions with potential assembly issues.

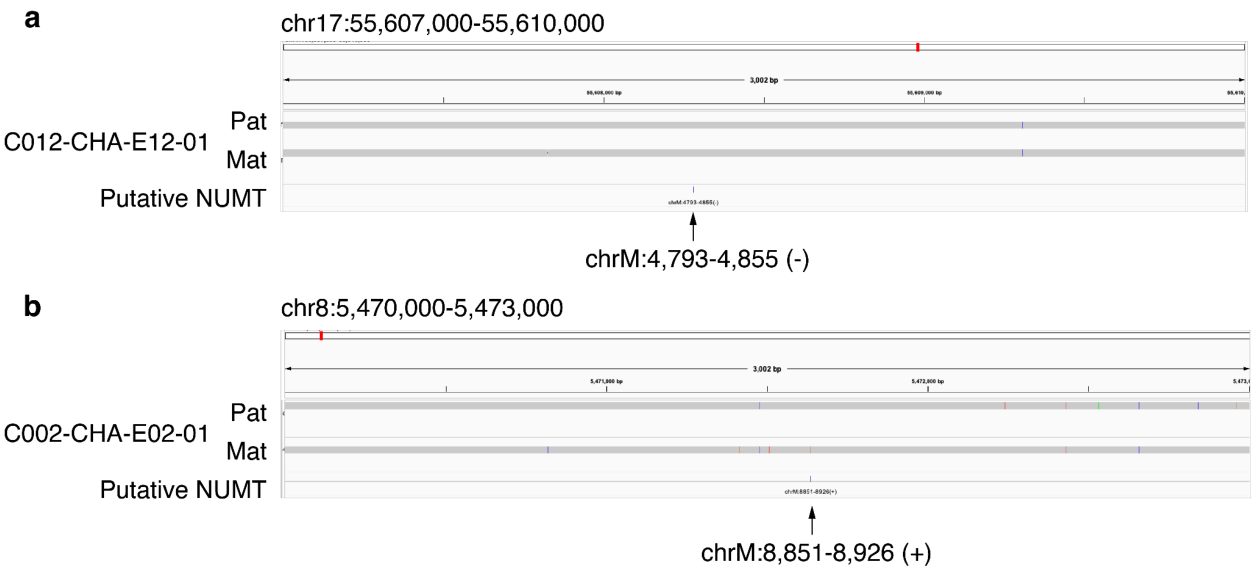


**Supplementary Figure 3.** Two NUMTs misidentified by short-read. No structural variants (SVs) or indels were detected between the individual genomes and the reference genome T2T-CHM13 within the NUMT regions identified by short-read sequencing, suggesting that these discrepancies result from mapping errors caused by short reads. The arrows indicate the nuclear sites and their corresponding mitochondrial regions misidentified by the short-read approach.

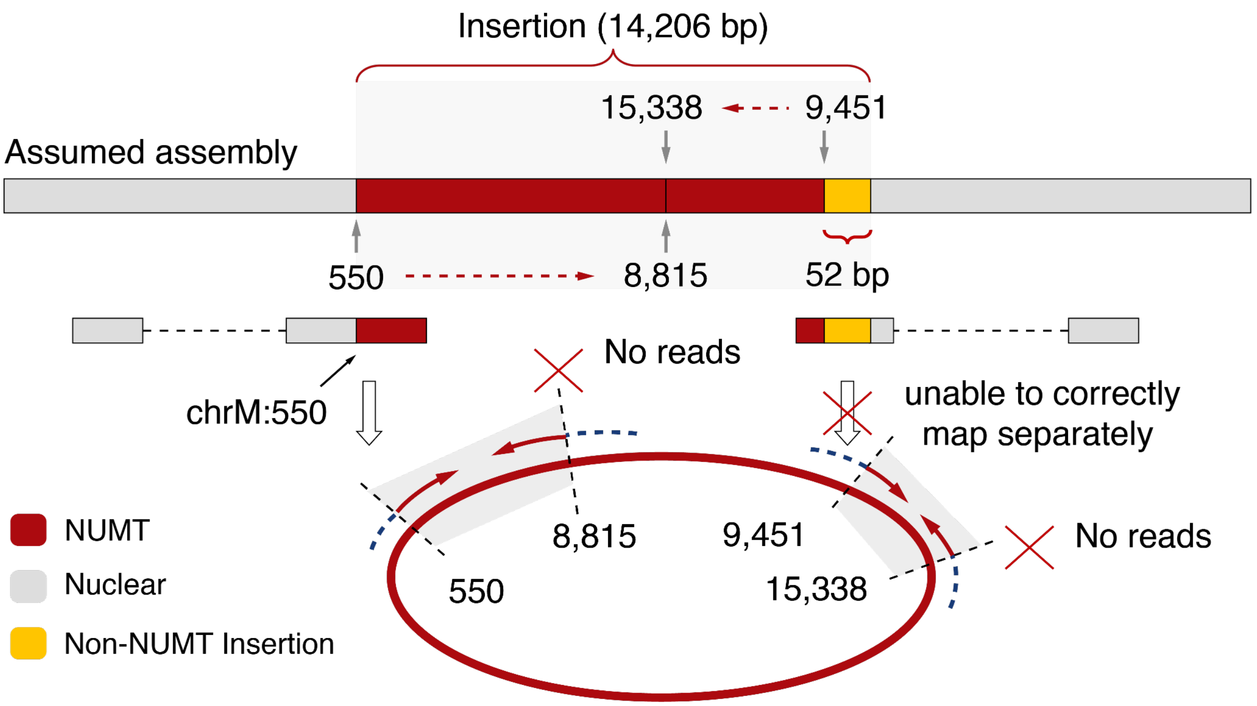


**Supplementary Figure 4.** A NUMT detected exclusively by PG-NUMT at chr6: 124,435,809-124,435,809 in T2T-CHM13. The NUMT comprises two distinct mtDNA-derived segments (chrM:550-8,815 and chrM:9,451-15,338). Reads spanning the right nuclear breakpoint cannot be accurately mapped or clipped due to a 52-bp insertion sequence adjacent to the breakpoint. Consequently, the precise structural organization of this NUMT remains incompletely resolved, as the absence of informative spanning short reads precludes definitive determination of its integration boundaries and internal structural configuration.

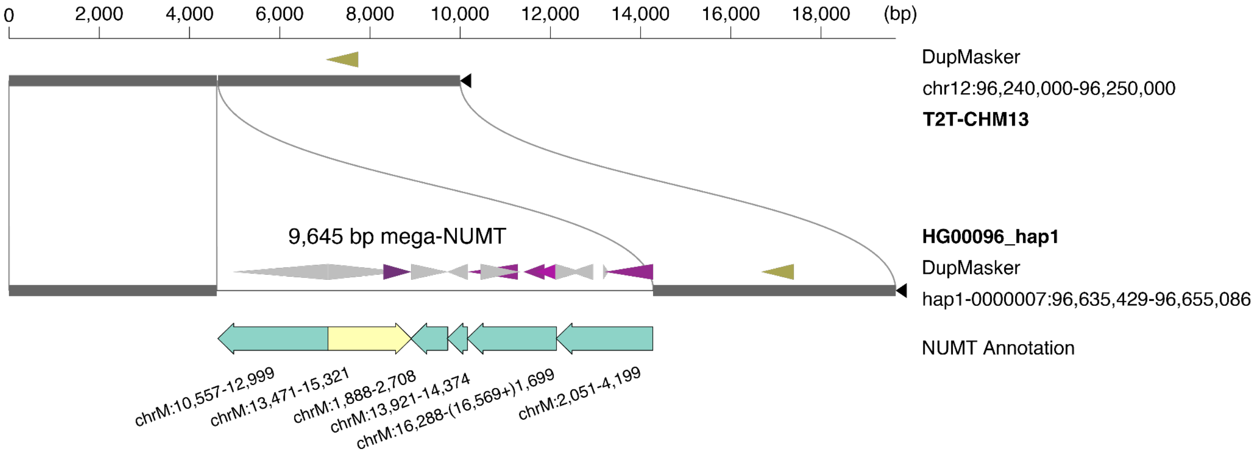


**Supplementary Figure 5.** A 9,645-bp mega-NUMT identified in this study comprises 6 distinct mtDNA-derived segments. The DupMasker annotation is attached to each genome segment.

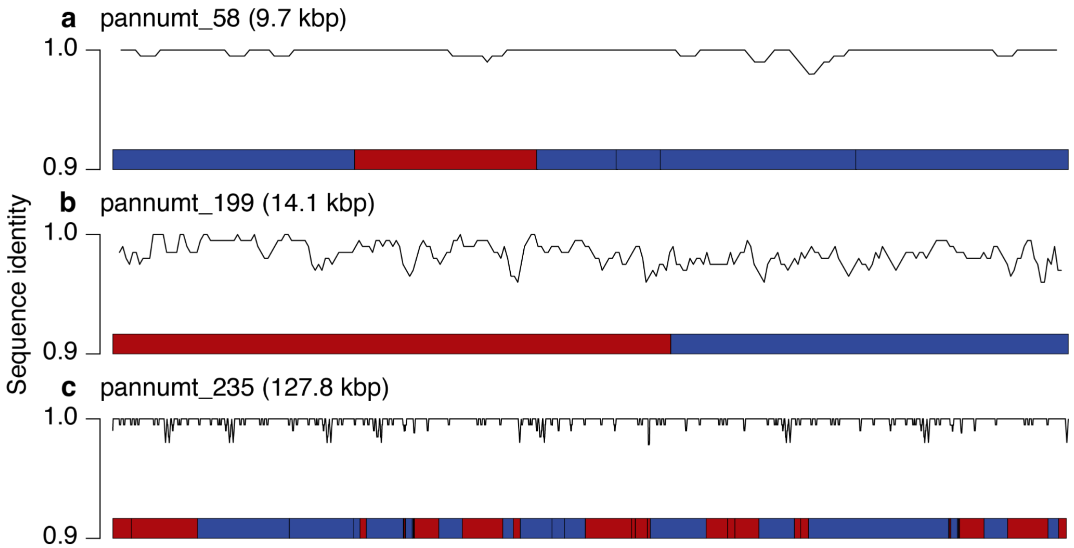


**Supplementary Figure 6.** Sequence identity patterns and structural organization of three mega-NUMTs (length ≥ 300 bp). The top panel displays the sequence identity of distinct segments relative to mtDNA. The bottom panel illustrates the schematic structure of the concatenated NUMT segments, where red and blue regions denote forward and reverse mtDNA strands, respectively.

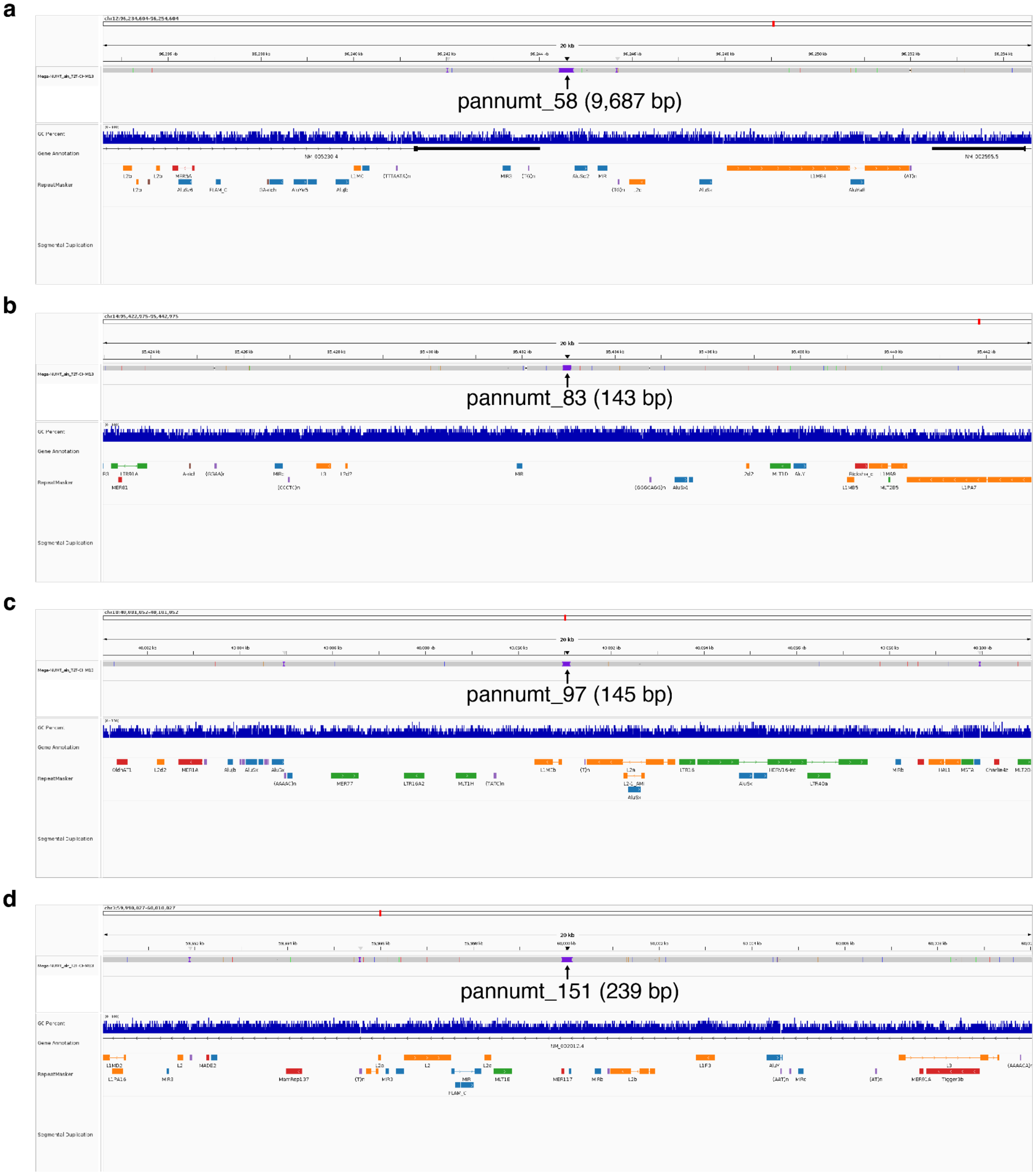
a
pannumt_58 (9,687 bp)
b
pannumt_83 (143 bp)
c
pannumt_97 (145 bp)
d
pannumt_151 (239 bp)

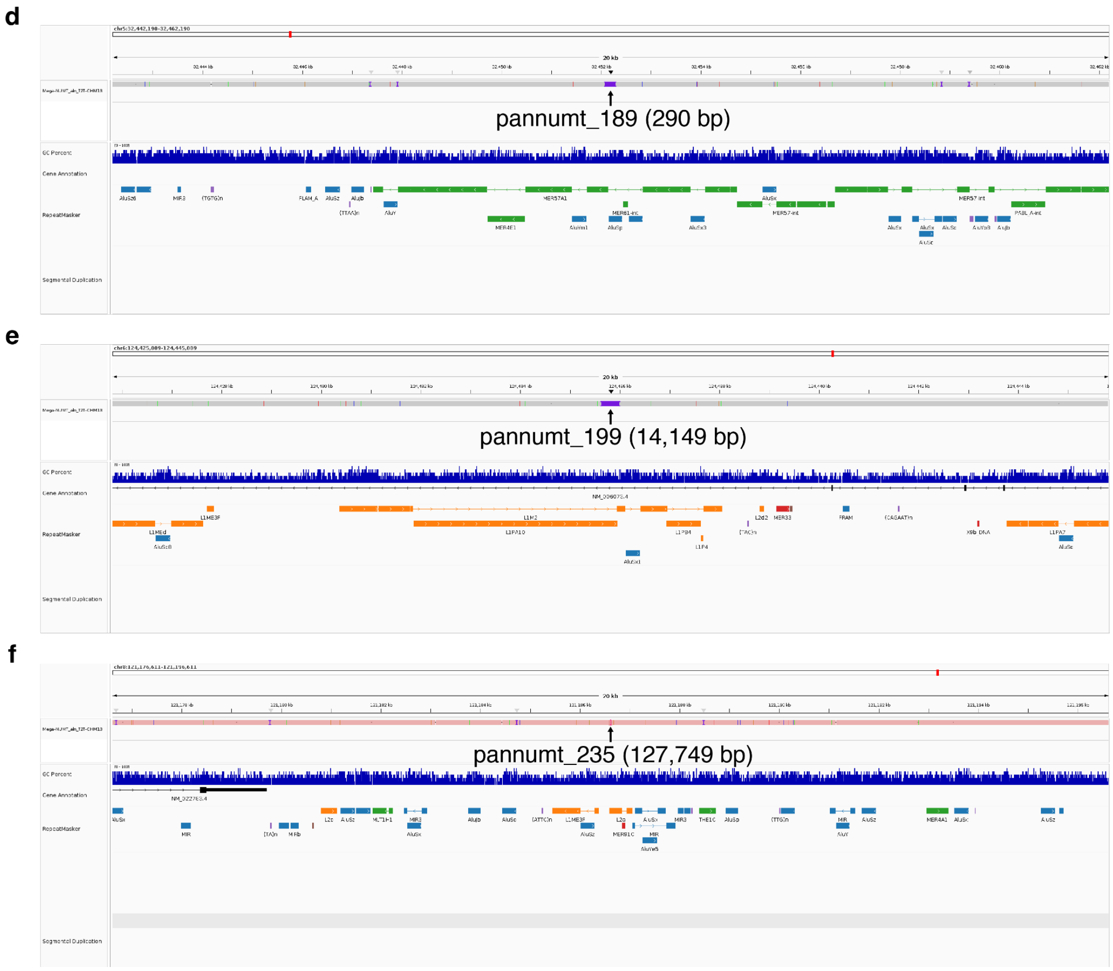


**Supplementary Figure 7.** Genomic context of seven mega-NUMTs. IGV screenshot showing the NUMT insertion and its genomic context. The top track illustrates the haplotype-to-reference alignment confirming the insertion. Subsequent tracks display (from top to bottom): GC content, gene annotations, repetitive elements (RepeatMasker), and segmental duplications.

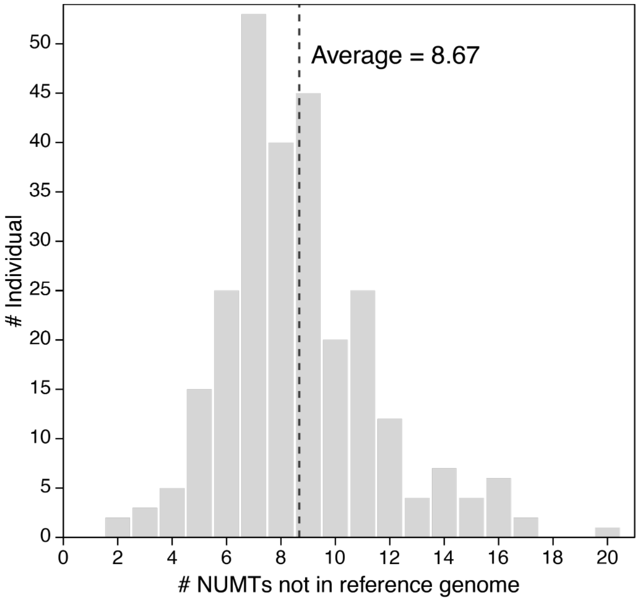


**Supplementary Figure 8.** The average number of NUMTs per individual that is not present in the reference genome.

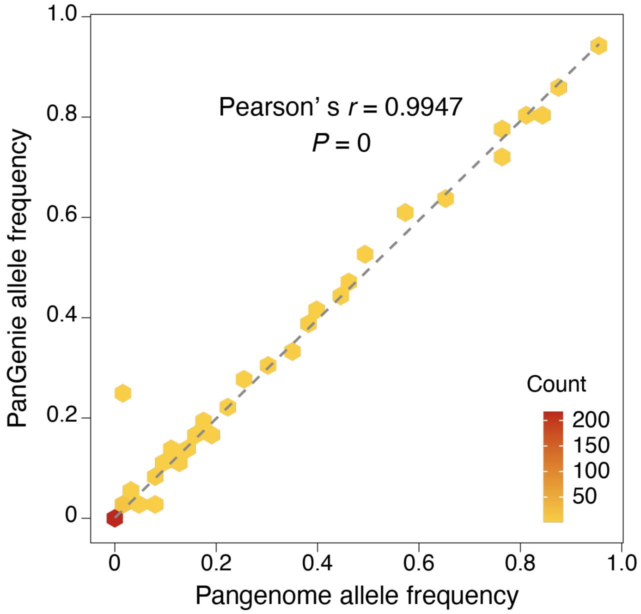


**Supplementary Figure 9.** Comparison of polymorphic NUMT allele frequencies between PanGenie genotypes and the pangenome graph across 40 HPRCp1 and 160 APGp1 individuals shows high concordance (Pearson's $r = 0.9947$, $P = 0$).

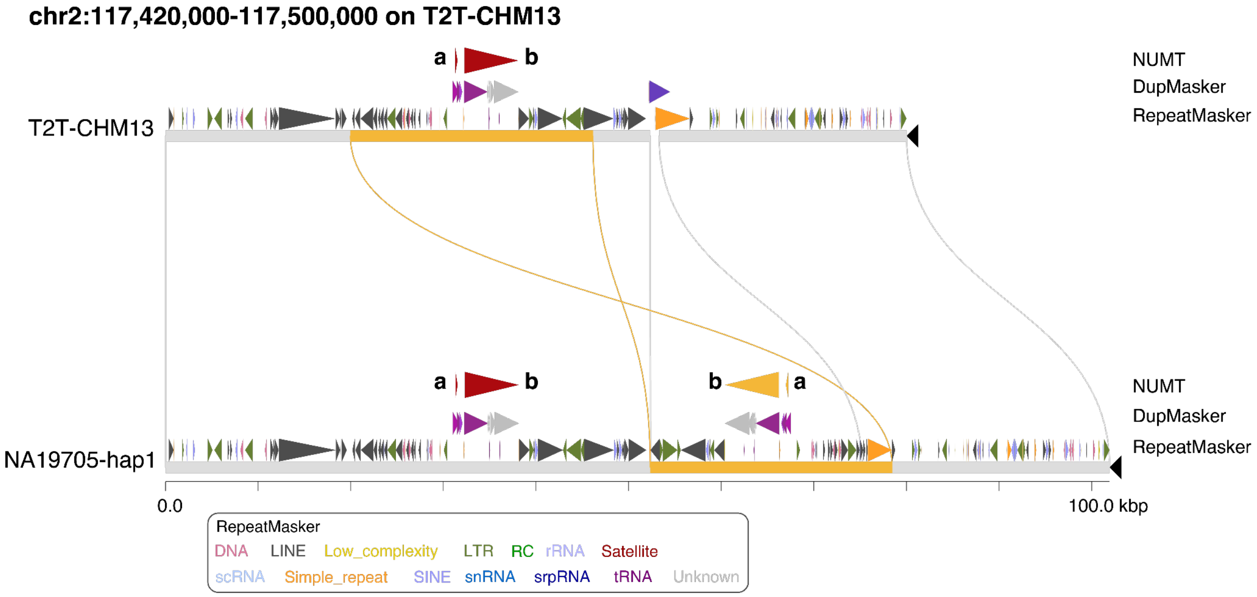


**Supplementary Figure 10.** The outlier locus of NUMT genotyping. This locus exhibits reduced genotyping accuracy due to adjacent repetitive sequences and the inversion structure. The DupMasker, RepeatMasker, and NUMT annotations are attached to each genome segment. Label a represents numt_384, while label b represents numt_385.

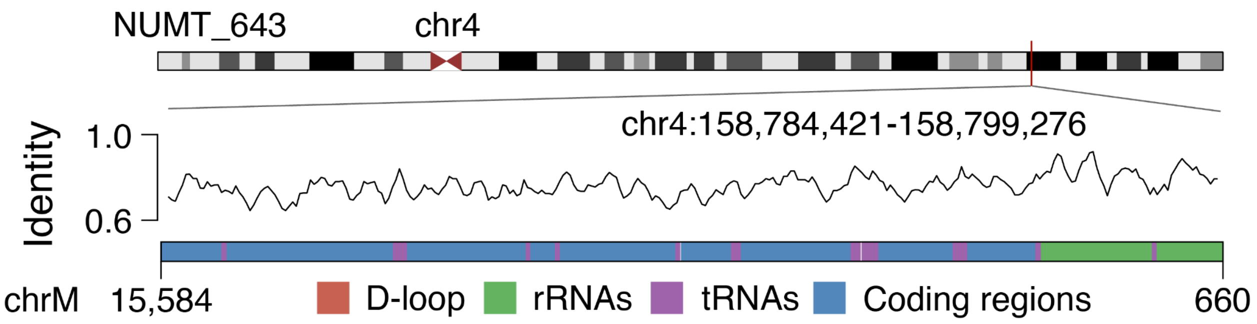


**Supplementary Figure 11.** The largest fixed NUMT at chr4:158,784,421-158,799,276 in T2T-CHM13. The sequence identity and the mtDNA annotation are shown at the bottom.

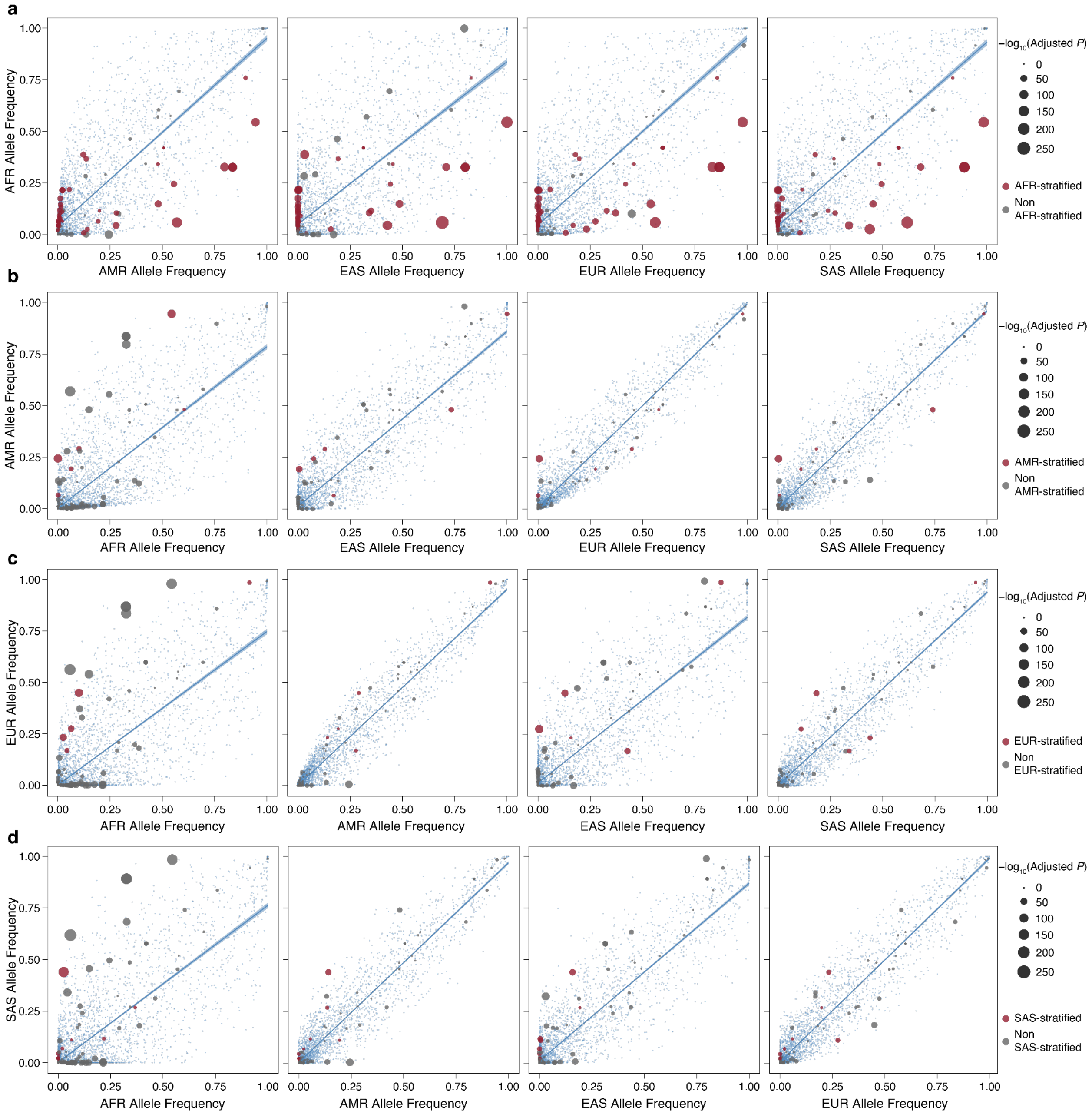


**Supplementary Figure 12.** Allele frequency comparison of NUMTs across superpopulations. Circles represent NUMTs, with those colored red indicating superpopulation-stratified NUMTs. Small dots denote SNV allele frequencies across superpopulations, serving as controls.

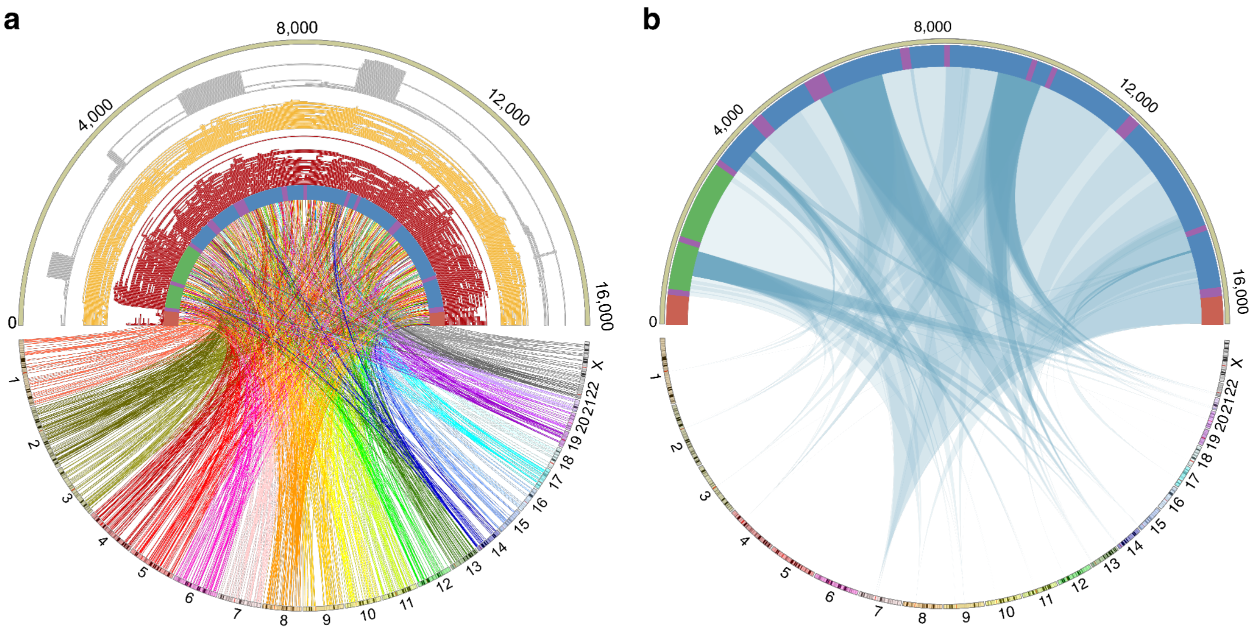


**Supplementary Figure 13.** The Circos plot of NUMT mtDNA origins. **(a)** The Circos plot illustrates the mtDNA origins of all NUMT types. The layers represent mtDNA annotation, fixed NUMTs, polymorphic NUMTs, and pericentromeric NUMTs from the inside to the outside. Links connect NUMT breakpoints between mtDNA and their corresponding nuclear genome insertion loci. **(b)** The Circos plot specifically illustrates the mtDNA origin of pericentromeric NUMTs. Links connect NUMT regions between mtDNA and their corresponding nuclear genome insertion regions.

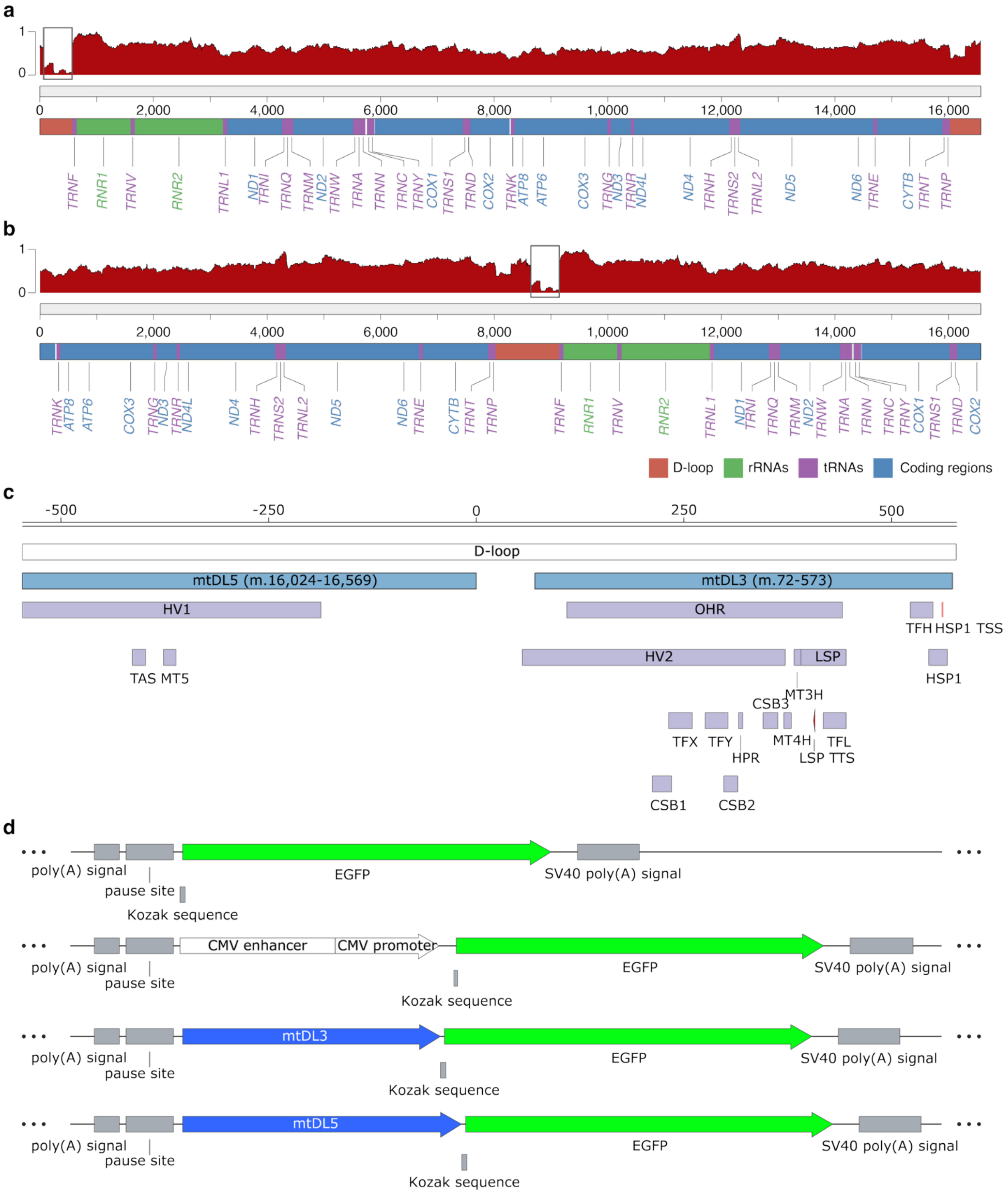


**Supplementary Figure 14.** Validation of fixed NUMT detection. The detection method employs two validation strategies: **(a)** using 1,000 distinct human mtDNA queries covering variants in the D-loop region across populations, and **(b)** using reference sequences with shifted start positions to avoid potential edge effects. Both results show decreased coverage in the 3′-end region of the

D-loop region and exclude alignment artifacts. The mtDL3 regions are highlighted with grey boxes. **(c)** The D-Loop structure annotation. mtDL5 corresponds to the mitochondrial DNA region chrM:16,024-16,569 and mtDL3 represents chrM:72-573. **(d)** Schematic diagram of plasmid constructs designed to assess the potential *cis*-regulatory activity of mtDL3-derived sequences. Both mtDL3 and mtDL5 were inserted into the plasmid backbone. The CMV enhancer and promoter were cloned into the same vector as the positive control, whereas the empty plasmid was used as the negative control.

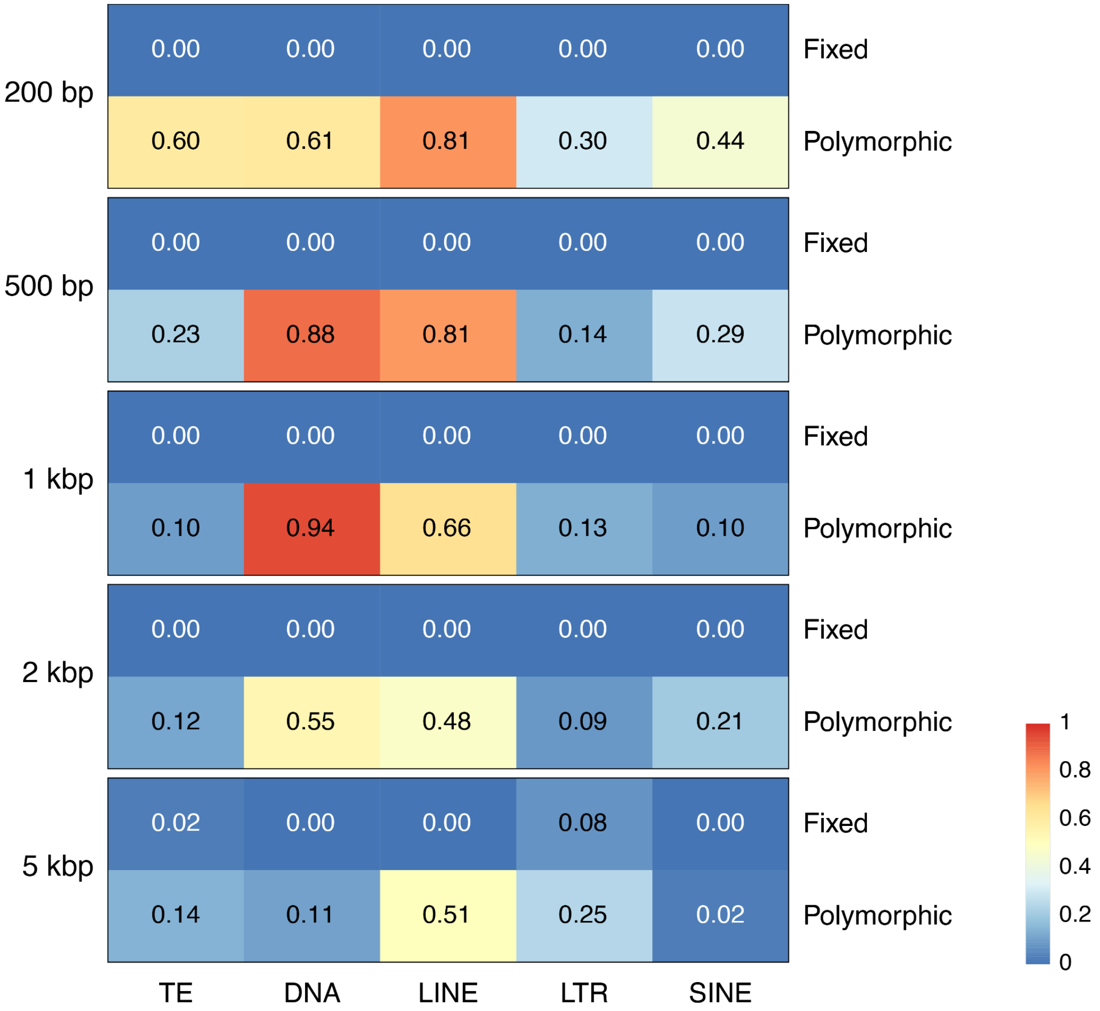


**Supplementary Figure 15.** Heatmap of enrichment analysis of fixed (top) and polymorphic (bottom) NUMT insertions associated with transposable elements in the human genome across different flanking window sizes. Values represent enrichment scores derived from permutation tests relative to random expectations (Methods). TE, transposable element; DNA, DNA transposon; LINE, long interspersed nuclear element; LTR, long terminal repeat; SINE, short interspersed nuclear element.

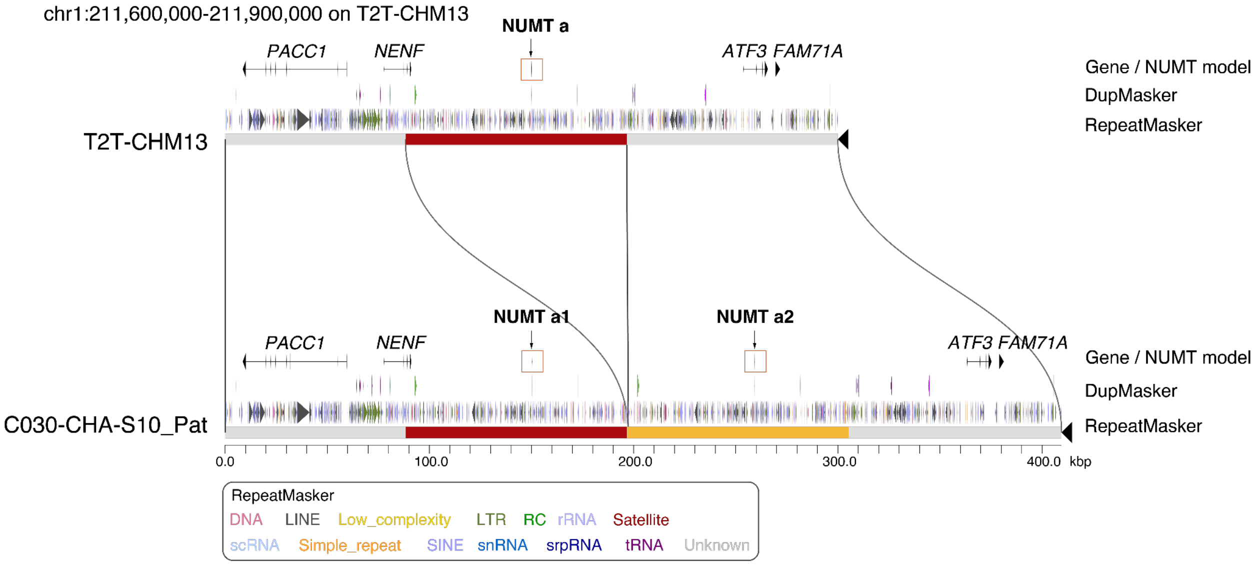


**Supplementary Figure 16.** NUMT formation resulted from intrachromosomal duplication. The genomic segment containing NUMTs is tandemly duplicated in the C030-CHA-S10_Pat, resulting in NUMT a1 and a2. NUMT a represents numt_47. The DupMasker, RepeatMasker, and NUMT annotations are attached to each genome segment.

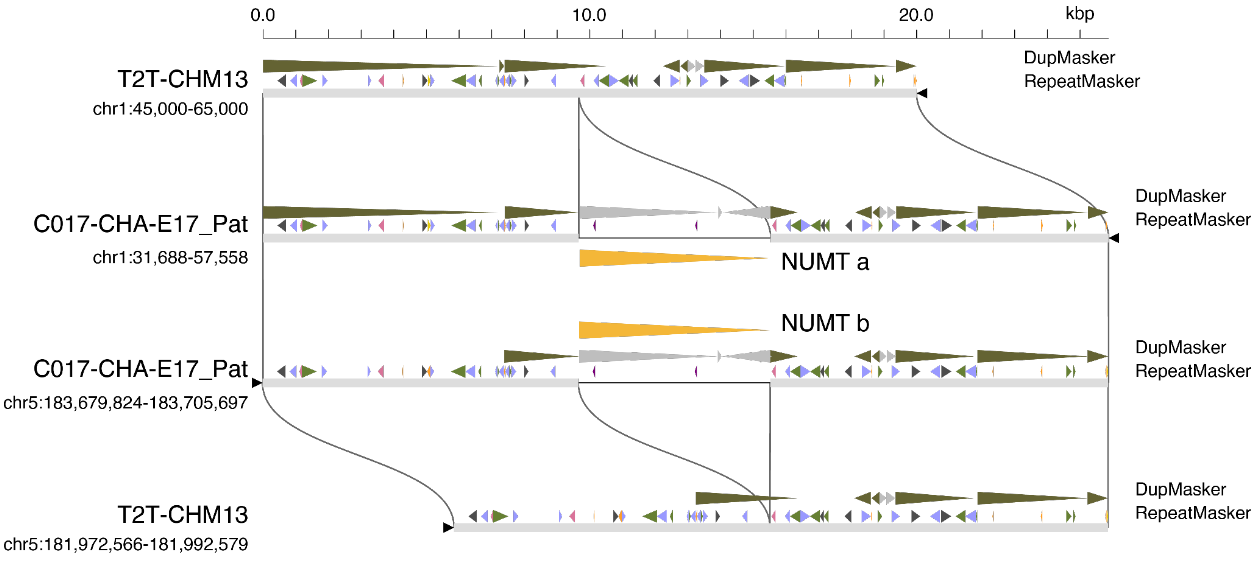


**Supplementary Figure 17.** NUMT formation resulted from interchromosomal duplication. NUMT a represents pannumt_18 and NUMT b represents pannumt_186. The DupMasker, RepeatMasker, and NUMT annotations are attached to each genome segment.

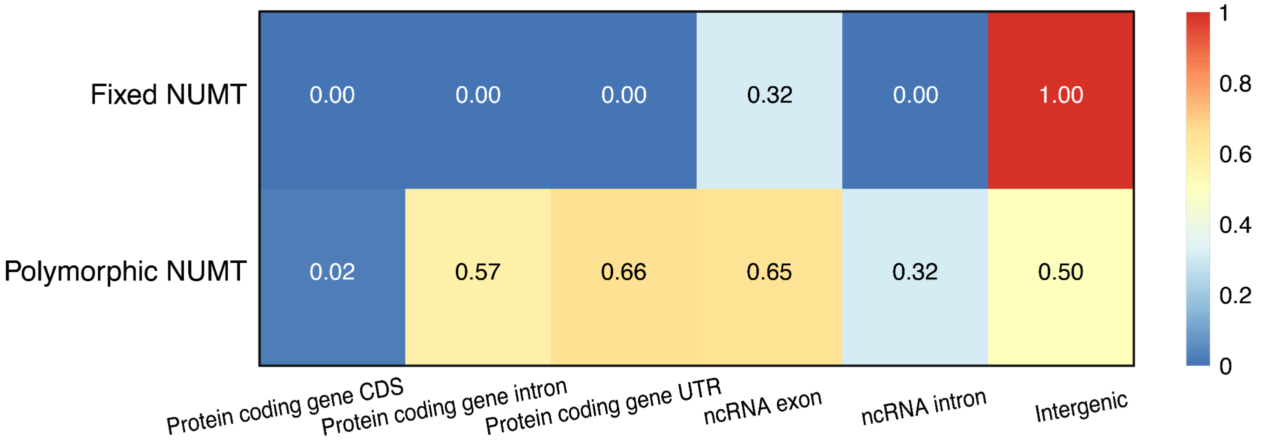


**Supplementary Figure 18.** Heatmap of enrichment analysis of fixed (top) and polymorphic (below) NUMT insertions across different genomic regions.

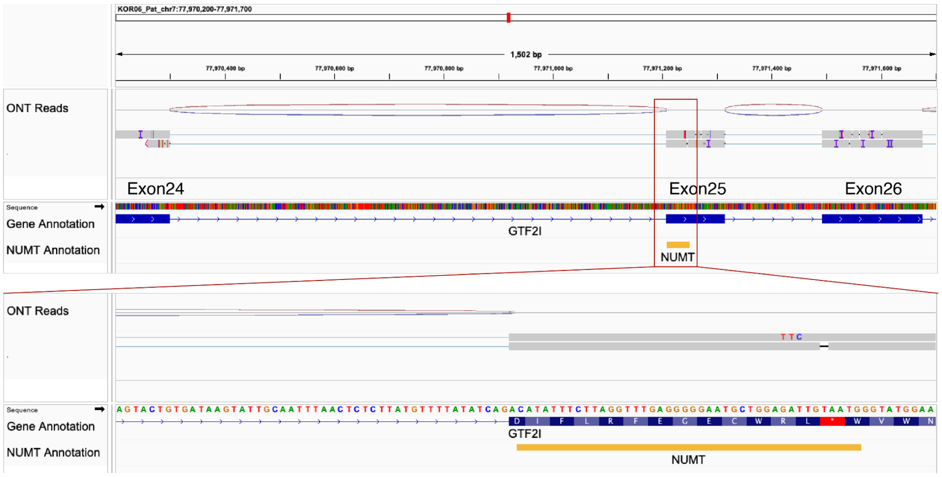


**Supplementary Figure 19.** Validation of the *GTF2I* NUMT insertion using Nanopore long-read RNA-seq. The top panel displays the alignment of RNA-seq reads to the *GTF2I* gene region in the carrier (KOR06-Pat), with the red box marking the 42-bp NUMT insertion in exon 25. The bottom panel provides a magnified view of the sequence and predicted translation at the insertion site. The insertion results in a frameshift mutation, introducing a premature termination codon (red asterisk *).

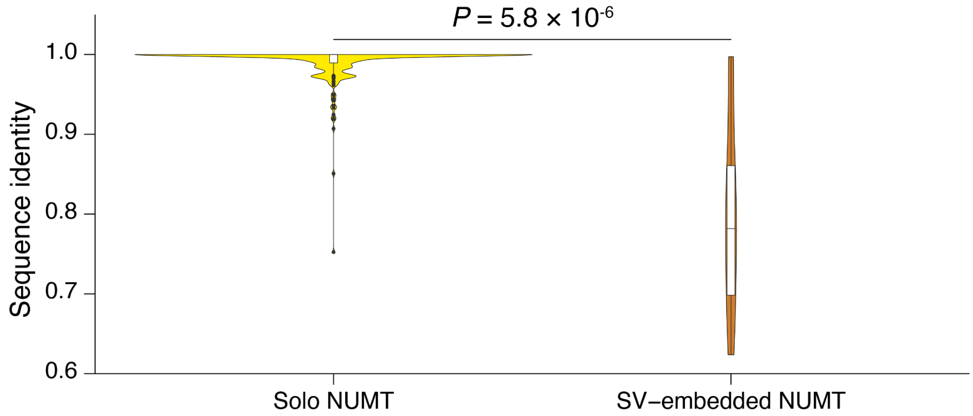


**Supplementary Figure 20.** Comparison of sequence identity to mtDNA between solo and SV-embedded NUMTs.

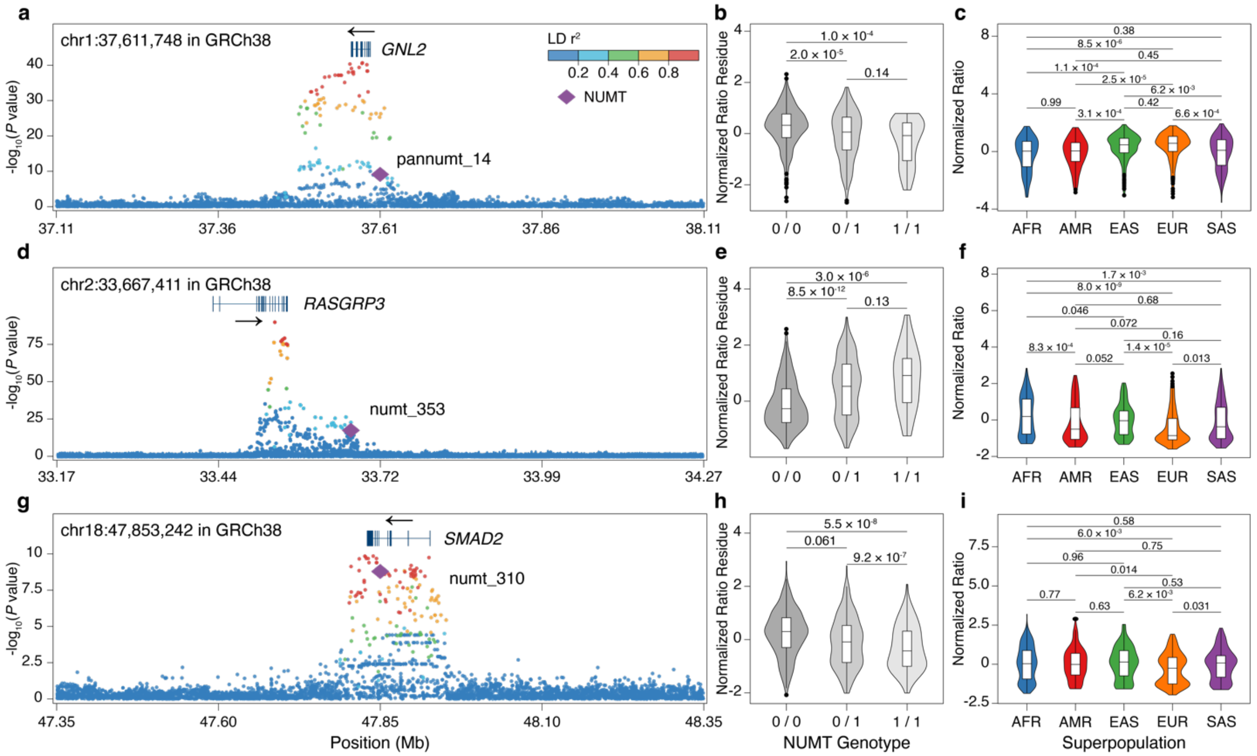


**Supplementary Figure 21.** The schematic representation of three NUMTs with dual QTL effects shows sQTL-associated splicing loci with peak statistical signals (lowest *P* values). The three sNUMT-sGene pairs include: pannumt_14 and *GNL2*_chr1:37,582,936-37,583,867 (a, b, c), numt_353 and *RASGRP3*_chr2:33,539,323-33,539,459 (d, e, f), numt_310 and *SMAD2*_chr18:47,883,964-47,896,521 (g, h, i).

**(a, d, g)** LocusZoom plots of regional sQTL association analyses of three NUMTs.

**(b, e, h)** Covariate-adjusted normalized intron excision ratio residue comparisons across NUMT genotypes.

**(c, f, i)** Normalized intron excision ratio comparisons among superpopulations.

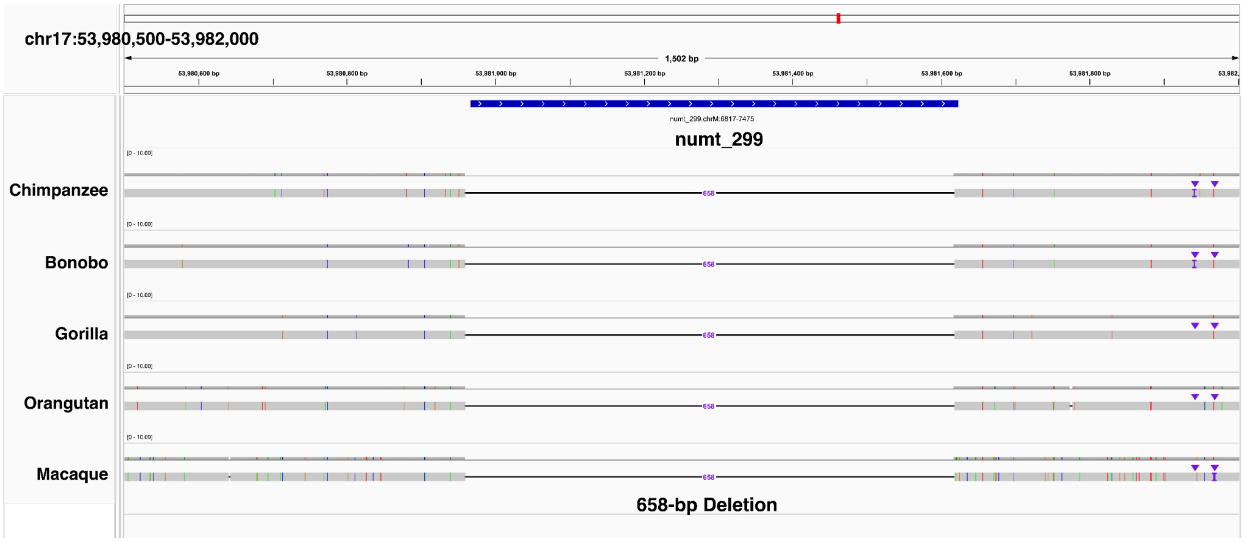


**Supplementary Figure 22.** An example of human-specific NUMT. The IGV tracks show the alignments of human individuals, apes, and macaques to T2T-CHM13v2.0, indicating that the insertion on numt_299 is a human-specific event.

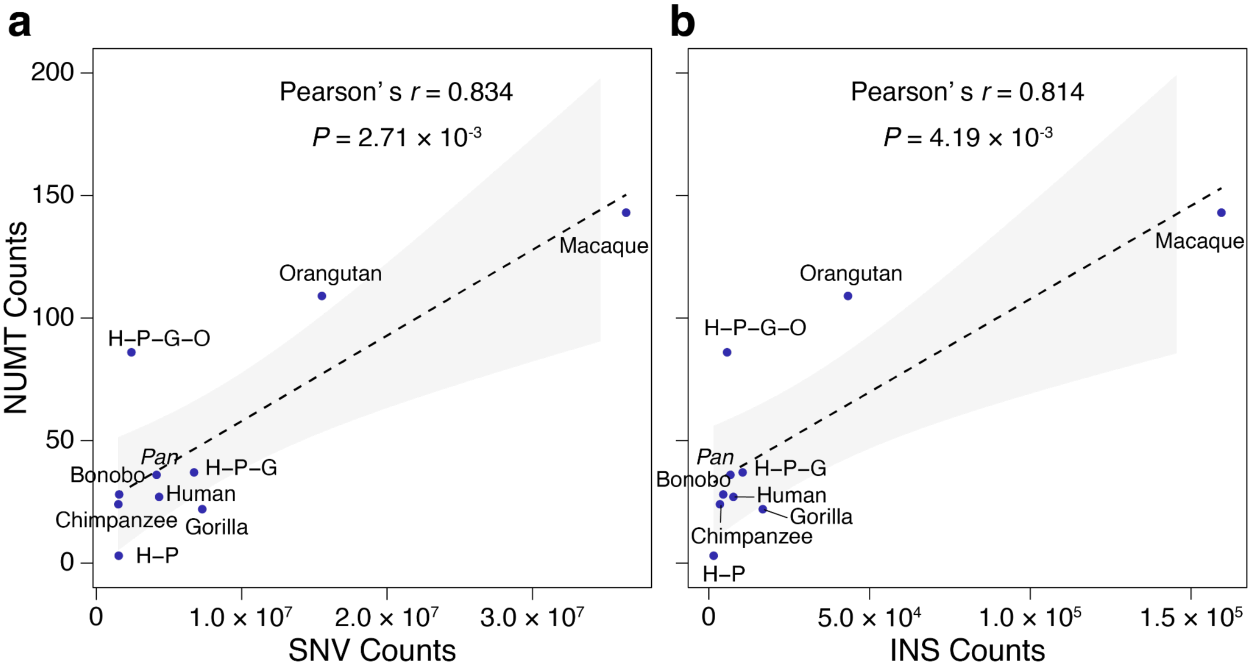


**Supplementary Figure 23.** The correlation between lineage-specific NUMT insertions and the accumulation of SNVs (a) and fixed insertions (b) in each lineage.

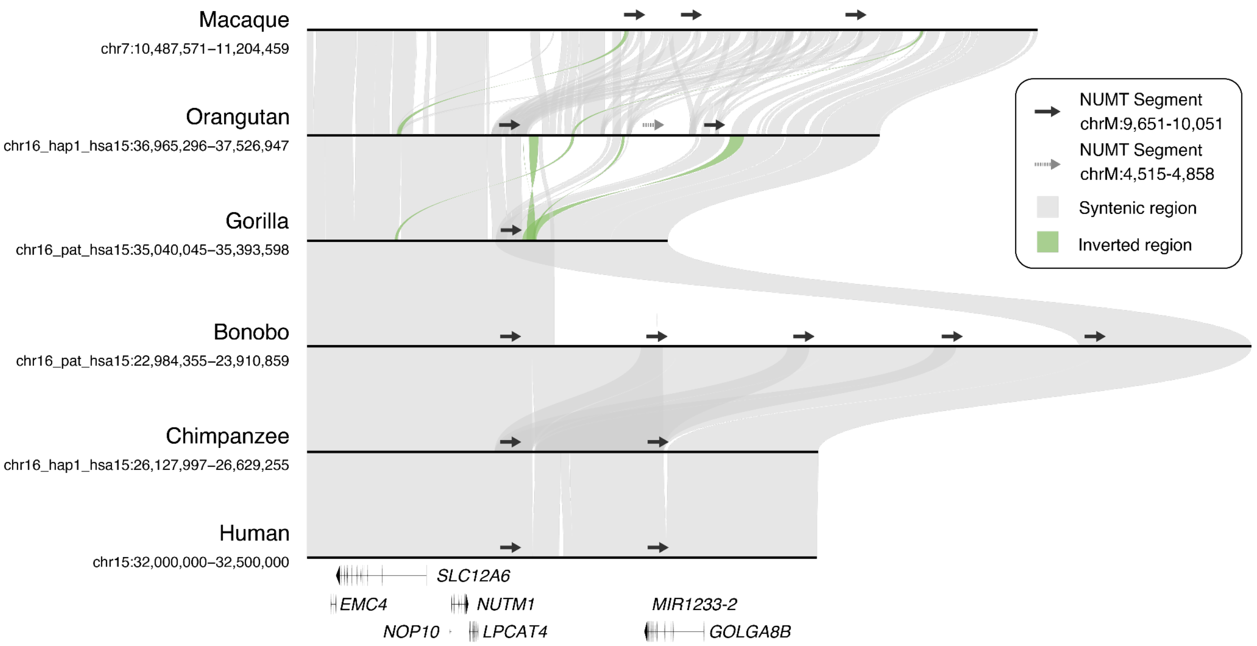


**Supplementary Figure 24.** Syntenic comparison of recurrent NUMTs across primates at chr15:32,000,000-32,500,000 in T2T-CHM13. The gene models are present at the bottom.

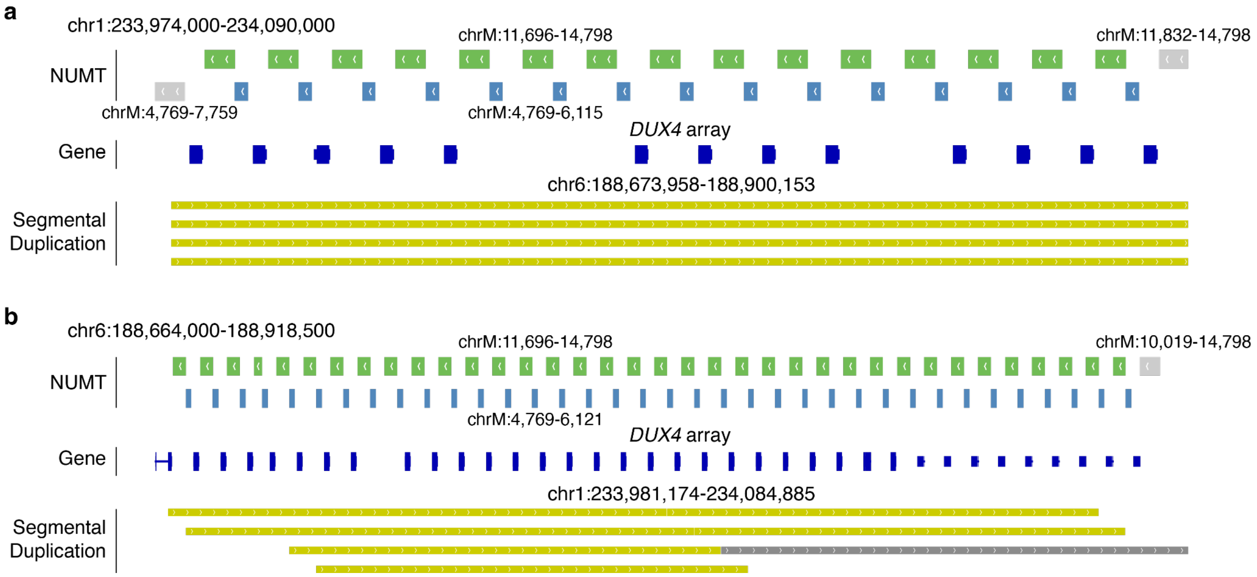


**Supplementary Figure 25.** Genomic organization of two large NUMT arrays in the macaque genome (T2T-MFA8). Tracks are arranged from top to bottom: NUMT annotations, gene annotations, and segmental duplications (SDs). In the NUMT annotation track, green blocks represent repeating units derived from chrM:11,696-14,798, while blue blocks denote units derived from chrM:4,769-6,115 (chr1 array) and chrM:4,769-6,121 (chr6 array). Grey blocks represent other NUMT segments with specific mtDNA coordinates. In the gene annotation track, blocks represent *DUX4* or *DUX4L* genes. The segmental duplication track indicates that the chr1 and chr6 arrays constitute segmental duplication pairs.

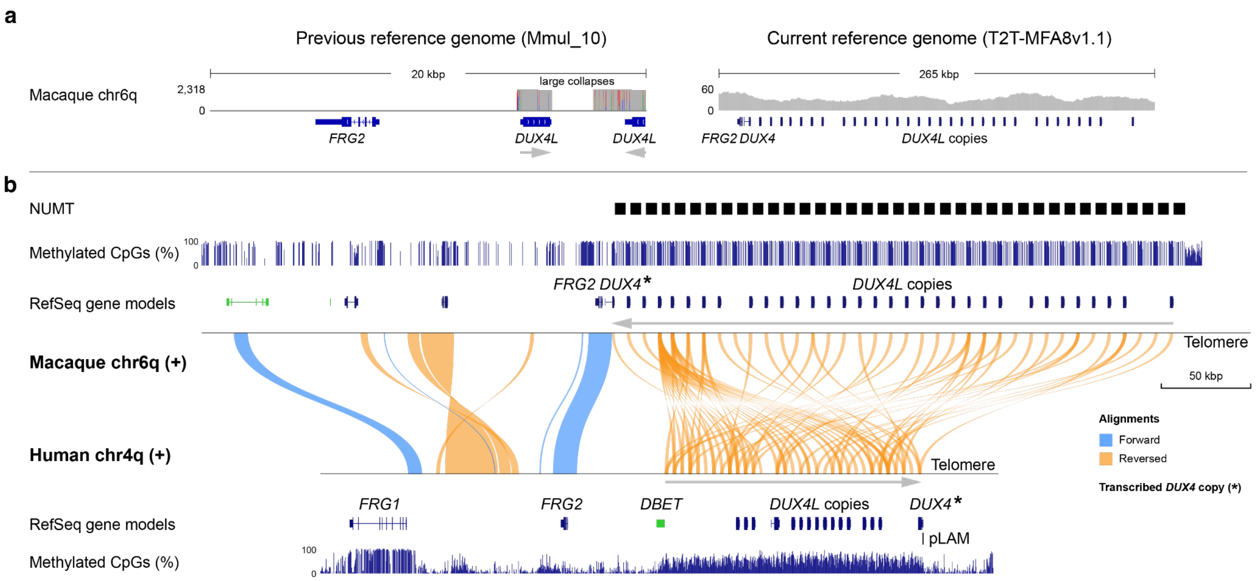


**Supplementary Figure 26.** The lineage-specific *DUX4*-NUMT array in macaque chr6. **(a)** Comparison of the *DUX4* locus between the previous reference (Mmul_10) and the current T2T assembly (T2T-MFA8), illustrating the successful resolution of previously collapsed large segmental duplications. **(b)** Comparative synteny and methylation landscapes of macaque (chr6q) and human (chr4q) *DUX4* arrays. The macaque array features *DUX4* and *DUX4L* copies interspersed with NUMT insertions (black boxes). CpG methylation tracks (blue bars) reveal that these NUMT insertions fragment the high-methylation region in macaques, in contrast to the continuous high-methylation profile observed in humans. Additionally, *DUX4* copies are oriented towards the telomere in humans with the most telomeric copy being intact, whereas the macaque arrays exhibit an inverted orientation. Moreover, a specific pLAM sequence acts as a permissive element in the disease mechanism of human Facioscapulohumeral Muscular Dystrophy (FSHD). Such pLAM is absent in macaques.

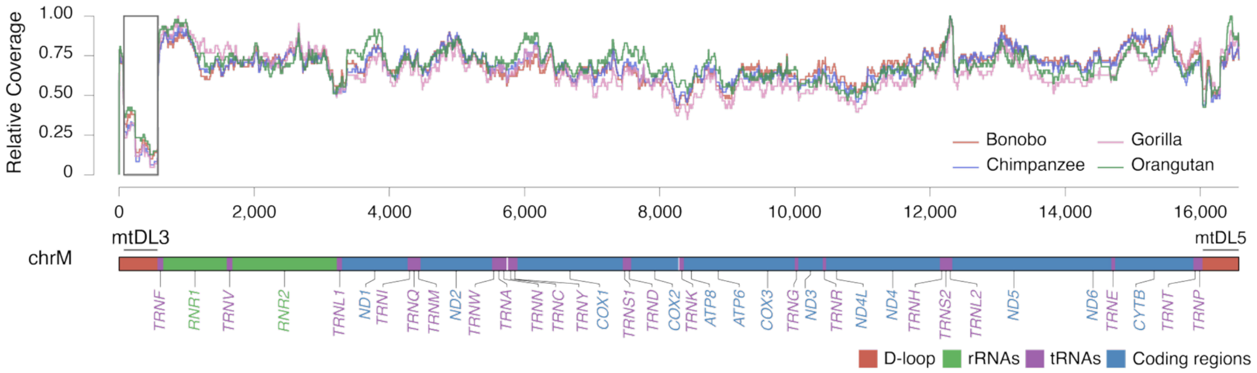


**Supplementary Figure 27.** Fixed NUMT coverage relative to the mtDNA coordinates across non-human primates. The x-axis represents the nucleotide positions of mtDNA, while the y-axis indicates the relative coverage distribution across fixed NUMT in distinct species (normalized by their maximum coverages). The mitochondrial gene annotation is presented at the bottom. The mtDL3 region (chrM:72-573) is highlighted with the grey box.

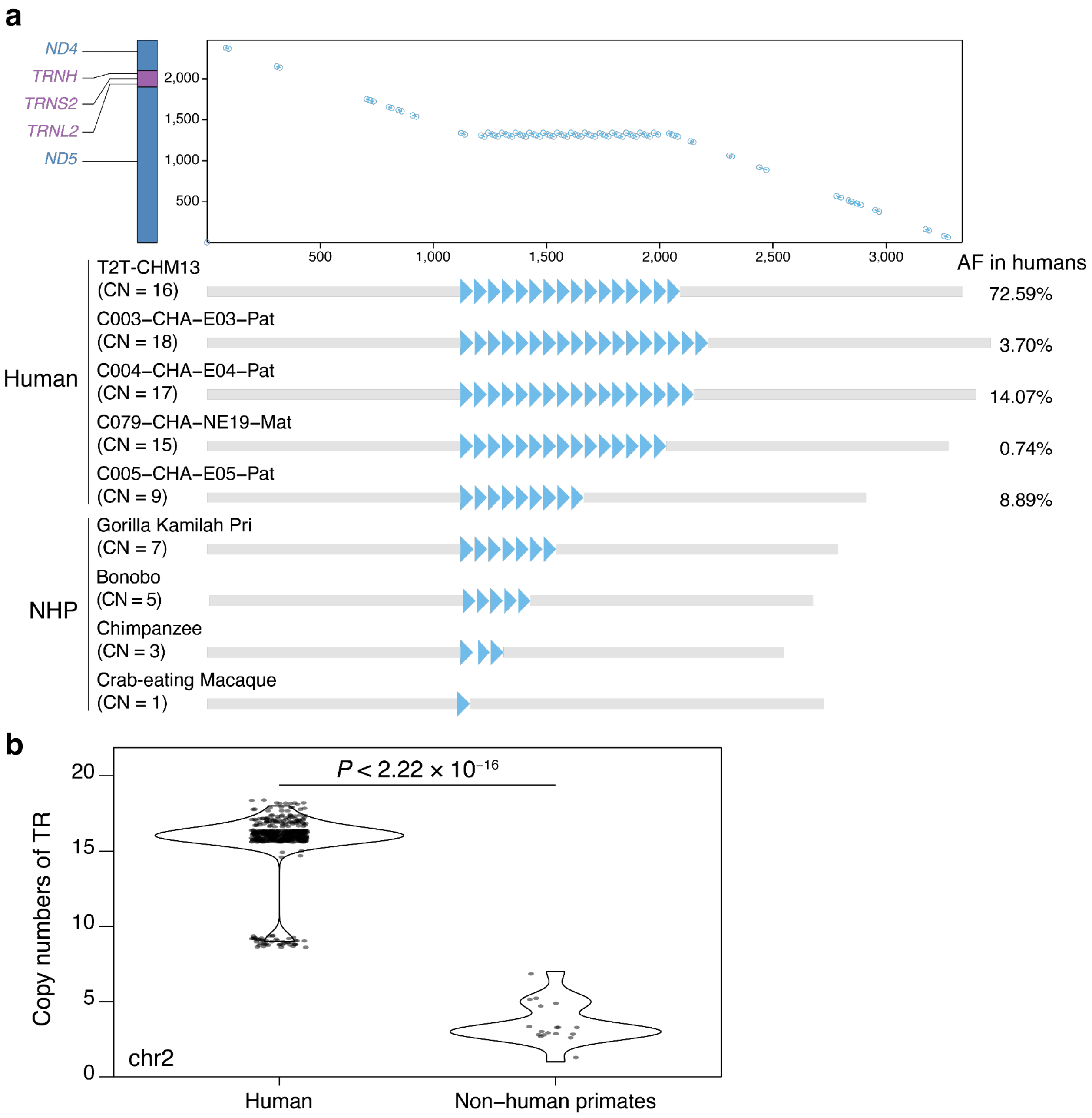


**Supplementary Figure 28.** A NUMT-derived VNTR expanded in humans (chr2:238,011,642-238,015,272 in T2T-CHM13).

**(a)** The syntenic comparison of the NUMT and mtDNA sequences is shown at the top. Schematic representation of the primate-specific NUMT-derived VNTR and its allele frequency distribution across humans is displayed at the bottom.

**(b)** Copy number of the tandem repeat motif across primates of the NUMT-derived VNTR.

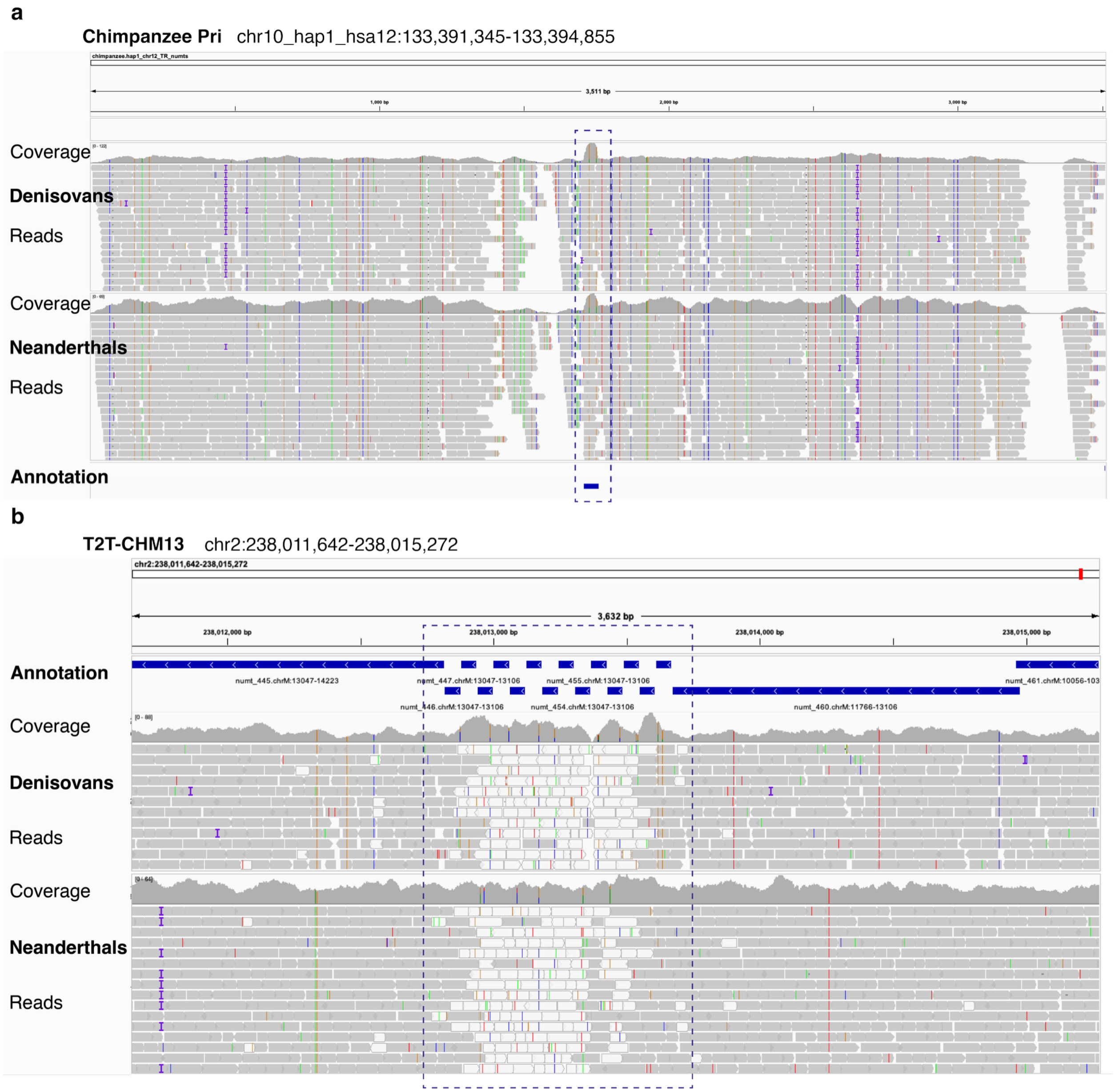


**Supplementary Figure 29.** Two NUMT-derived VNTRs in archaic humans, including locus chr12:126,591,633-126,592,357 (a) and locus chr2:238,011,642-238,015,272 (b). Coverage of archaic human short-reads mapped to chimpanzee (a) and human (b) reference genomes shows no dip in NUMT-derived VNTR regions, indicating multi-copy TRs in Neanderthals and Denisovans. NUMT-derived VNTR regions are highlighted with blue boxes.

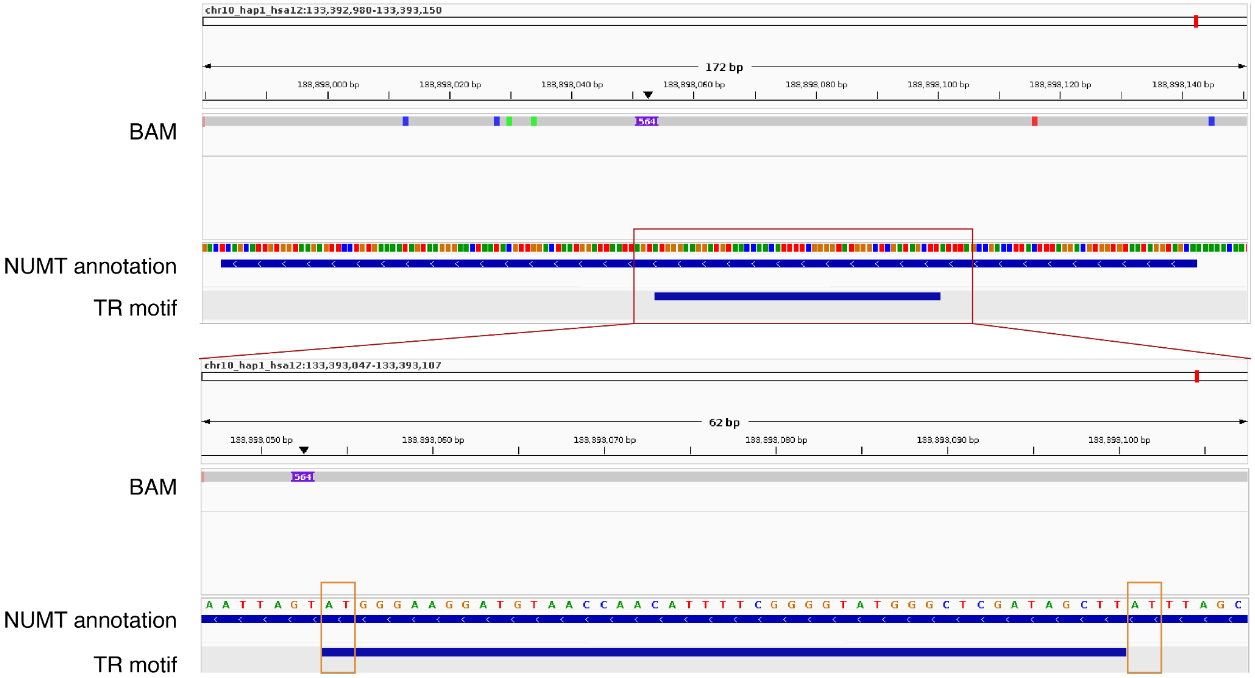


**Supplementary Figure 30.** Evidence of microhomology-mediated expansion in a human-specific NUMT-derived VNTR (chr12 in human and chr10 in chimpanzee). The top panel displays (from top to bottom): the alignment of the human genome (query) to the chimpanzee genome (ref), the chimpanzee NUMT annotation, and the consensus repeat motif annotation in the chimpanzee. The bottom panel provides a magnified view of the repeat unit boundaries. Orange boxes highlight the 2-bp microhomology (“AT”) identified at the 5’ and 3’ boundaries of the repeat unit, providing the structural basis for replication slippage and subsequent VNTR array expansion.

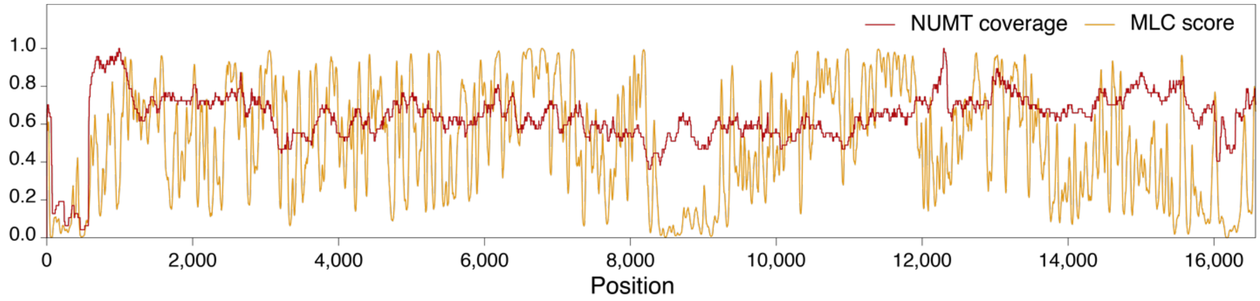


**Supplementary Figure 31.** Comparison of fixed NUMT coverage and mtDNA Local Constraint (MLC) Score. The x-axis represents nucleotide positions of mtDNA, while the y-axis indicates the relative coverage distribution of fixed NUMTs (normalized to their maximum coverages; red) and the MLC score (orange). Both NUMT coverage and MLC score exhibit low values in the mtDL3 region.
